# Ultra-fast physics-based modeling of the elephant trunk

**DOI:** 10.1101/2024.10.27.620533

**Authors:** Bartosz Kaczmarski, Derek E. Moulton, Zéphyr Goriely, Alain Goriely, Ellen Kuhl

## Abstract

With more than 90,000 muscle fascicles, the elephant trunk is a complex biological structure and the largest known muscular hydrostat. It achieves an unprecedented control through intricately orchestrated contractions of a wide variety of muscle architectures. Fascinated by the elephant trunk’s unique performance, scientists of all disciplines are studying its anatomy, function, and mechanics, and use it as an inspiration for biomimetic soft robots. Yet, to date, there is no precise mapping between microstructural muscular activity and macrostructural trunk motion, and our understanding of the elephant trunk remains incomplete. Specifically, no model of the elephant trunk employs formal physics-based arguments that account for its complex muscular architecture, while preserving low computational cost, to enable fast screening of its configuration space. Here we create a reduced-order model of the elephant trunk that can–within a fraction of a second– predict the trunk’s motion as a result of its muscular activity. To ensure reliable results in the finite deformation regime, we integrate first principles of continuum mechanics and the theory of morphoelasticity for fibrillar activation. We employ dimensional reduction to represent the trunk as an active slender structure, which results in closed-form expressions for its curvatures and extension as functions of muscle activation and anatomy. We create a high-resolution digital representation of the trunk from magnetic resonance images to quantify the effects of different muscle groups. We propose a general solution method for the inverse motion problem and apply it to extract the muscular activations of three representative trunk motions: picking a fruit; lifting a log; and lifting a log asymmetrically. For each task, we identify key features in the muscle activation profiles. Our results suggest that, upon reaching maximum contraction in select muscle groups, the elephant trunk autonomously reorganizes muscle activation to perform specific tasks. Our study provides a complete quantitative characterization of the fundamental science behind elephant trunk biomechanics, with potential applications in the material science of flexible structures, the design of soft robots, and the creation of flexible prosthesis and assist devices.

## 1. Motivation

The intricate musculature of the elephant trunk has fascinated scientists and engineers for decades. Along with other biological structures such as the octopus arm or the human tongue, it is a muscular hydrostat in which the near-incompressible muscle tissue enables various motion mechanisms without the support of a rigid skeleton [1, 2, 3]. Carefully orchestrated contractions of around 90,000 muscle fascicles of the elephant trunk [4] can generate a wide variety of complex motion archetypes. Figure 1 illustrates the spectrum of motion, ranging from the precise manipulation of small objects [5, 6] to the high force generation to lift heavy objects [7].

**Figure 1:**
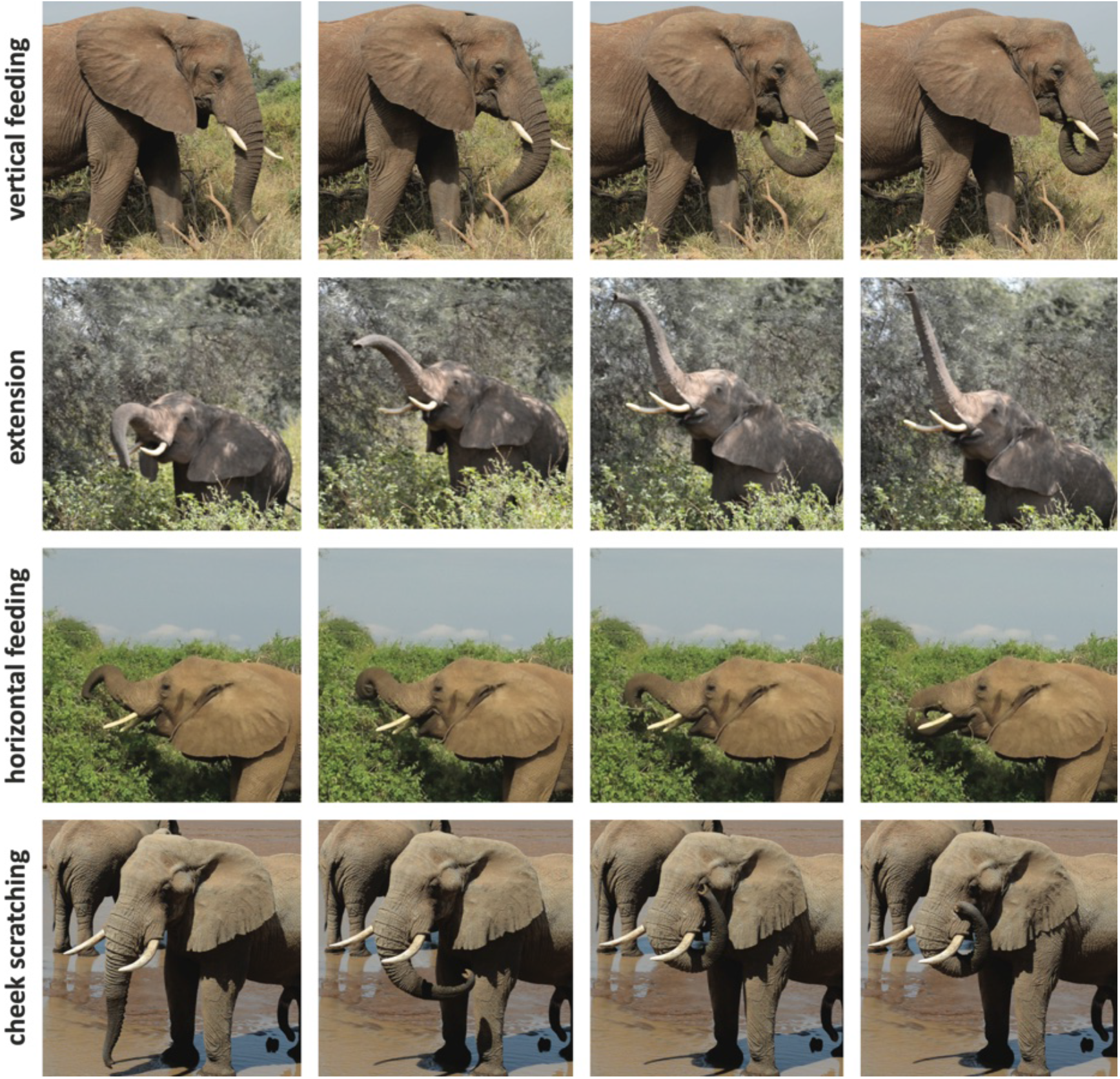
The elephant trunk can perform complex motions. These series of natural uses performed by African elephants (*Loxodonta africana*) were recorded in the Samburu National Reserve in Kenya, Summer 2024.

From the perspective of biomechanics, the elephant trunk is an example of a soft slender structure, in which the activation of the internal muscles determines its deformation in a tight interplay with applied external loading and boundary conditions. However, while the mechanics of passive slender structures have seen centuries of general theoretical treatise [8, 9, 10, 11, 12, 13, 14, 15, 16], system-specific modeling [17, 18, 19, 20, 21, 22, 23, 24, 25], and experimentation [26, 27, 28, 29, 30, 31, 32, 33], studies of slender structures with internal mechanical activation remain sparse and require further improvements in computational efficiency and fidelity [34, 35, 36, 37, 33].

In particular, the elephant trunk, despite being one of the most prominent examples of active slender structures in nature, remains incompletely understood and is still a subject of active research. Past work involved experimental studies [38, 4, 5, 39, 40, 41, 42] and modeling efforts [43, 44, 45, 46, 47] which provided invaluable insights into the principles that underlie the elephant trunk’s control capabilities and investigated the main features of the trunk that engineers could extract to produce more effective designs in soft-robotic applications. In fact, the sheer versatility of the elephant trunk served as an inspiration for a large range of engineering solutions developed throughout the years [48, 49, 50, 51, 52]. Additionally, other muscular hydrostats, such as the octopus arm, have sparked the development of computational tools that have deepened our understanding of the immense mechanical complexities inherent in this class of structures [53].

However, to our knowledge, there is no model of the biomechanics of the elephant trunk that derives from first principles and finite-deformation continuum mechanics to ensure reliable simulation results. Proposed models use over-simplifying assumptions in the mechanical formulation or are computationally expensive which prevents fast solution of inverse problems and efficient exploration of the trunk’s configuration space. In this work, we fill these research gaps by providing a reduced-order physics-based model of the elephant trunk’s musculature and use its real-time implementation to simulate the biomechanics of the elephant trunk for several motion tasks. While reproducing trunk motions in a simulation setting is generally challenging, we are able to use our digital trunk representation to automatically extract the muscular activations underlying the physiological trunk motions. As a result, the model not only provides key quantitative insights into the mechanical principles governing slender biological systems, but also provides guidance for future development of engineered active structures.

We begin by introducing the general concept of tapered active slender structures [54] and define their deformation as a function of the internal fibrillar activation and the applied external loading. We then specialize the general model to construct a representation of the elephant trunk consisting of multiple muscular subdomains with various muscle architecture types. We derive the trunk curvature and extension formulas in response to fibrillar activation and simplify the result for the case of uniform activation and linear tapering. By using magnetic resonance images of the elephant trunk [38], we extract the geometrical and architectural parameters of the trunk muscles, resulting in 28 independently activated muscular subdomains. Further, we incorporate the effects of trunk incompressibility, optimize the computational structure of the model implementation to enable real-time simulation, and define the kinematic principles of the motion of the trunk’s proximal base. Using the derived model, we establish general biomechanical principles governing the trunk representation by analyzing the signs and magnitudes of the contributions of different muscle groups to the overall trunk motion. Finally, we construct and analyze three elephant trunk motions through optimization of muscle fiber contractions.

## 2. Elephant trunk mechanics

### Reduced-order continuum model

We model the elephant trunk as an active slender structure, in which an internal mechanical activation induced by the muscular contractions contributes to its deformation in addition to any external loading. Here we describe the continuum mechanics of the trunk according to the reduced-order theory of active slender structure mechanics [54], as shown in fig. 2.

**Figure 2:**
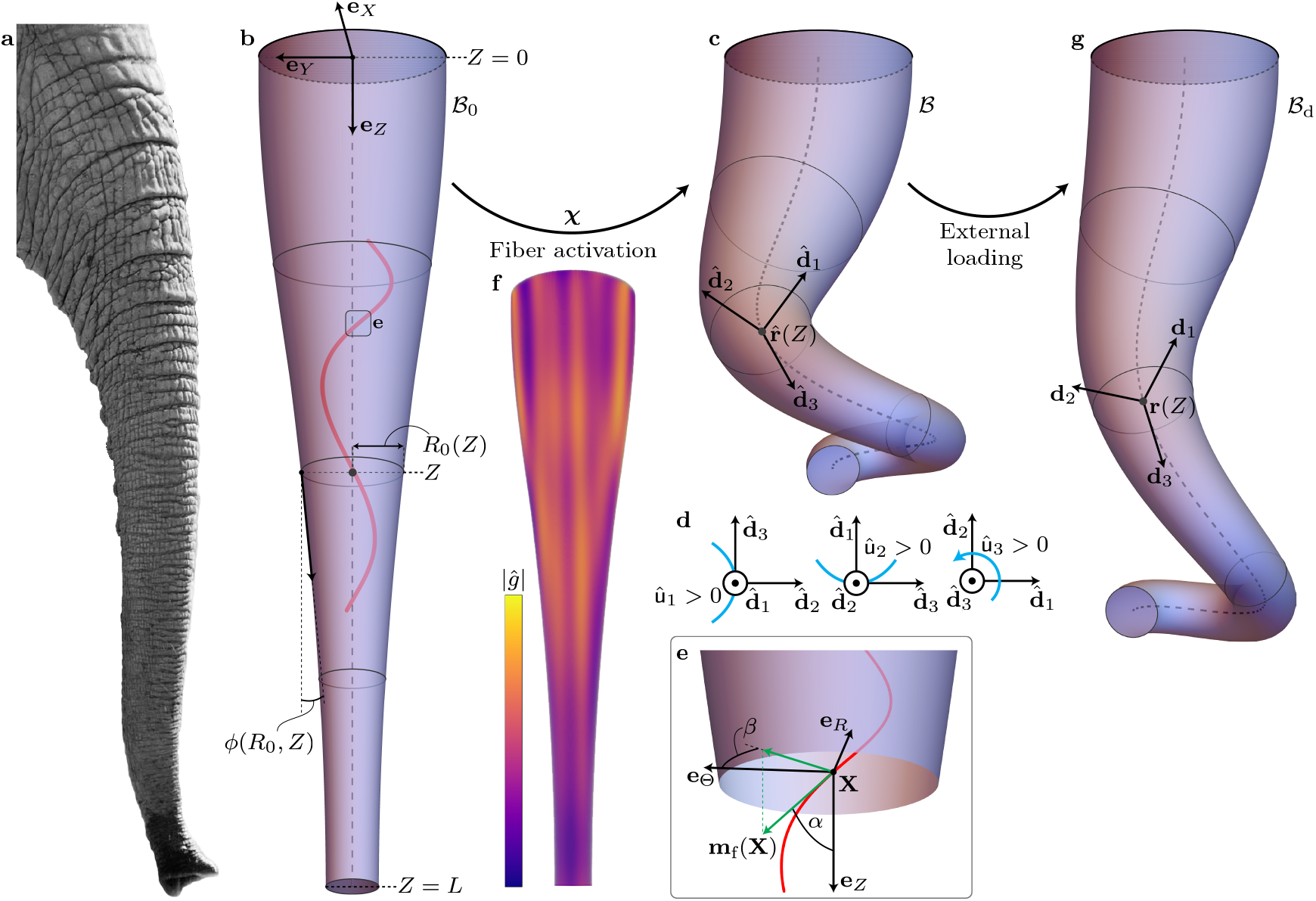
Reduced-order continuum model of the elephant trunk as a tapered active slender structure. (a) The elephant trunk. Internal muscular activation governs the deformation of the trunk in addition to the external loading. Image adapted from Pexels. (b) The initial configuration **ℬ**_0_ representing the trunk prior to fibrillar activation and external loading. We define the domain geometry with *Z* ∈ [0, *L*] and an outer radius profile *R*_0_(*Z*), assuming disk cross sections along **e**_*Z*_ at all *Z*. (c) The activated configuration **ℬ** resulting from a fiber activation field *ĝ* applied over a fiber direction field **m**_f_. The deformation *χ* uses a one-dimensional representation of **ℬ** in terms of the activated centerline 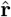 and director basis 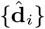.(d) Sign convention for the bending components û_1_, û_2_, and the twist density component û_3_ of **û**. (e) Definition of the fiber direction field **m**_f_. At a given point **X, m**_f_(**X**) makes an angle *a*(**X**) with **e**_*Z*_ and an angle *β*(**X**) with **e**_Θ_ in the plane spanned by **e**_*R*_ and **e**_Θ_. (f) Example of an arbitrary scalar field *ĝ*(**X**) applied throughout the initial configuration. The field |*ĝ*(**X**)| represents the magnitude of muscular activation at all points **X** in **ℬ**_0_. (g) The deformed configuration **ℬ**_d_ defined by the centerline **r** and directors {**d**_*i*_} that result from an applied external loading.

We begin by defining the tubular initial configuration **ℬ**_0_ of the trunk in a cylindrical basis {**e**_R_, **e**_Θ_, **e**_*Z*_} with respect to the cylindrical coordinates {*R*, Θ, *Z*}, as well as the activated configuration **ℬ** in a basis {**e**_*r*_, **e**_*θ*_, **e**_*Z*_} with coordinates {*r*, *θ*, *z*}. We assume that every cross section along **e**_*Z*_ of the structure **ℬ**_0_ is a disk with a slowly varying radius *R*_0_(*Z*), for *Z* ∈ [0, *L*], where *L* is the total length of the structure. We then use dimensional reduction to reduce the computational complexity of the model and enable purely analytical tractability of the deformation solution. Specifically, we represent the deformation in terms of a one-dimensional centerline **r**(*Z*) that describes the trajectory traced by the slender configuration, and an orthonormal director basis {**d**_1_, **d**_2_, **d**_3_} that defines the orientation of the cross section at **r**(*Z*), for all *Z* ∈ [0, *L*]. The kinematics of the resulting one-dimensional filamentary structure dictate the following general relationships between the centerline and the director basis (see [55] for details on Kirchhoff rod formulation):

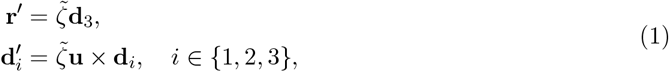

where the derivatives are with respect to 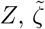,denotes the axial extension of the structure, and 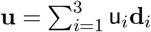 is the Darboux curvature vector [56]. For the deformation of the activated configuration **ℬ**, we denote the centerline by 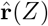 and the director basis by 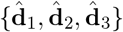giving the kinematic relations 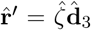 and 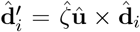,for *i* ∈ {1, 2, 3}, where 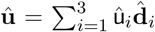 are the curvatures in the activated configuration.

Using the centerline and the directors, we can express the reduced deformation map χ : **ℬ**_0_ → **ℬ** as

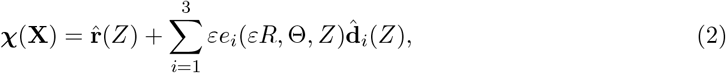

where **X** ∈ **ℬ**_0_ is a point in the initial configuration, *ε* is the ratio of the characteristic cross-sectional radius to the length *L*, and *εe*_*i*_ are the reactive strains that define the deformation in the cross section. The quantity *ε* is the small parameter of the model that permits the homogenization used in deriving the explicit deformation expressions in the theory [54, 57]. Figures 2b and c visualize the deformation χ in terms of the centerline 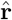 and the director basis 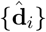 of the activated configuration. The functions 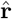 and 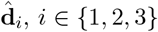,act as maps from the interval *Z* ∈ [0, *L*] to ℝ^3^. In fig. 2d, we specify the sign convention for the activated curvatures and twist density û_1_, û_2_, and û_3_.

We define the internal activation process representing the muscular contraction as a pre-strain contraction or elongation of internal fibers arbitrarily distributed throughout the continuum **ℬ**_0_ [54]. Most generally, we can write a vector field of fiber directions in the cylindrical basis as

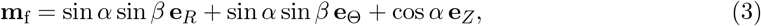

where *α*(**X**) ∈ (−*π*/2, *π*/2] and *β*(**X**) ∈ (−*π*/2, *π*/2] are the fiber direction angles at all points **X** ∈ **ℬ**_0_; see fig. 2e. The fiber direction field gives rise to the activation tensor **G** with eigenvectors **m**_f_, **m**_f, ⊥_ = cos *β* **e**_R_ — sin *β* **e**_Θ_, and 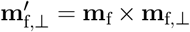 where the corresponding eigenvalues define the contractions or elongations due to activation along the three orthonormal eigenvectors. To define the deformation map **χ** in terms of the internal activation process, we decompose multiplicatively the deformation gradient **F** into the elastic part **A** and the defined activation part **G** , such that[55,58]

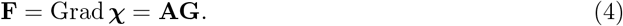

Using this decomposition of **F**, we then minimize the total energy of the system with respect to all cross sectional deformations admissible under **χ** for an arbitrary fiber direction field **m**_f_ and a general distribution of fibrillar activations throughout that field.

The minimization procedure, omitted here, yields the following closed-form formulas for the curvatures û_*i*_ and the extension 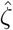 of the activated configuration (details given in [54]):

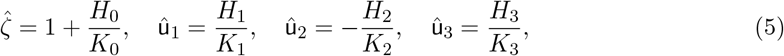

where

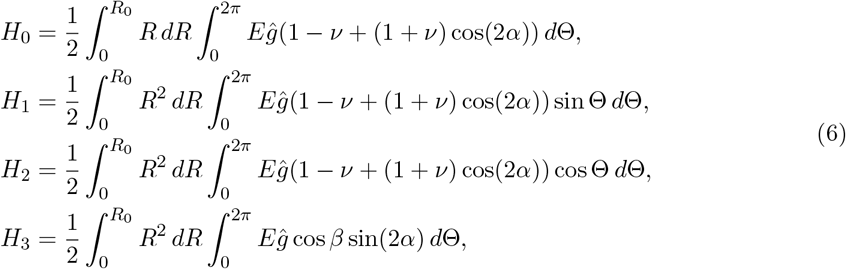

where *α* = *α*(*R*, Θ, *Z*) and *β* = *β*(*R*, Θ, *Z*) are the fiber architecture angle functions, *E* = *E*(*R*, Θ, *Z*) is the Young’s modulus, *ν* = *ν*(*R*, Θ, *Z*) is the Poisson’s ratio, and *ĝ* = *ĝ*(*R*, Θ, *Z*) is a scalar function defining the extent of fibrillar activation at each point in the domain. Figure 2f shows an example of an arbitrary fibrillar activation field *ĝ* applied throughout the initial configuration. The denominator quantities *K*_0_, *K*_1_, *K*_2_, and *K*_3_ are the stiffness coefficients, classically encountered in the theories of rod mechanics, which simplify to

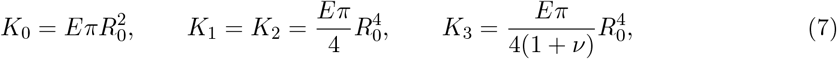

in the case of a disk cross section.

The musculature of the elephant trunk contains three primary muscle architectures: longitudinal, helical, and radial [4]. Under the assumption of disk cross sections, these architectures simplify the general *α*(*R*, Θ, *Z*) and *β*(*R*, Θ, *Z*) functions to angles *α* and *β* that are constant with respect to Θ in the whole domain. In the case of a constant outer radius *R*_0_(*Z*) = *R*_0_, *α* = *β* = 0 for longitudinal, *α* ∈ (−*π*/2, *π*/2) \ {0} and *β* = 0 for helical, and *α* = *β* = *π*/2 for radial fibers. However, since the elephant trunk is a tapered structure, the fiber architecture has to reflect the tapering of the outer radius *R*_0_(*Z*) along *Z*. According to [54], given a tapering angle function (*R, Z*) that is independent of Θ, we obtain the tapered fiber field by transforming a fiber direction

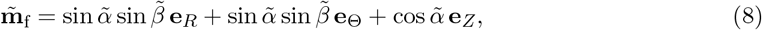

using a rotational transformation by *ϕ*(*R, Z*) about –**e**_Θ_, which gives the tapered fiber field

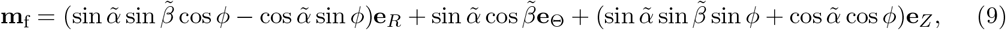

where 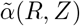 and 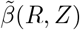 are auxiliary angle functions expressed in the pre-tapered space. We use 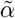 and 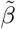 throughout this manuscript since their spatial variation is significantly easier to express compared to the physical angles *α* and *β*, which we can obtain from the relationships [54]

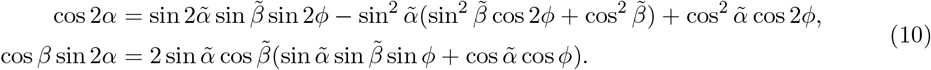

### External loading

Equations (5)-(7) define the intrinsic extension and curvatures generated by arbitrary fibrillar activations. To compute the deformation due to not only the internal muscular activation, but also the external loading, we define a deformed configuration **ℬ**_d_—the result of deforming the activated configuration **ℬ** through external forces. We assume that the motion of the elephant trunk is slow enough such that, at every time point, **ℬ**_d_ is a quasi-static solution of the force and moment balance equations for extensible Kirchhoff rods [56]:

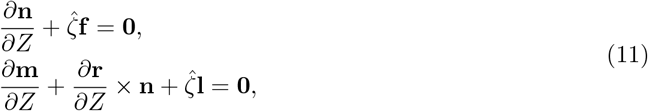

where **n** is the internal force, 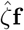 is the external body force per unit length of **ℬ**_0_, **m** is the internal moment, **r** is the centerline function of the deformed configuration, and 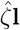 is the external body couple per unit length of **ℬ**_0_. We provide in fig. 2g a schematic of a sample deformed configuration **ℬ**_d_ generated through external loading applied to **ℬ**. To compute **ℬ**_d_, we solve the boundary value problem defined in eq. (11) with a choice of **f** , **l**, and boundary conditions on **m**(*Z*) and **n**(*Z*) appropriate for a given loading scenario.

Previous work posed the boundary value problem resulting from the weight of an active slender structure due to gravity [46, 59]. Here, we consider an extended library of loading scenarios consisting of:

- *A gravitational body force due to the weight of the trunk;*
- *A point load* **N** = ∑ N_*i*_**d**_*i*_ *applied at Z* = *L;*
- *A uniformly distributed load* **w**(*Z*) *given by*

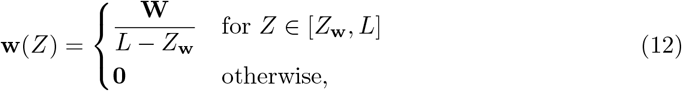

caused by an object of weight **W** ∈ ℝ^3^ lifted by the trunk, and applied along a one-dimensional contact region 𝒞 : [*Z*_**w**_, *L*] → **ℬ**;
- *A uniformly distributed body couble* 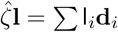 given by

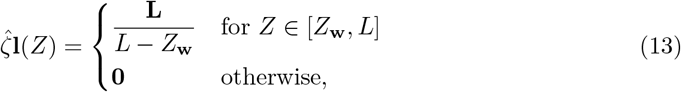

caused by a moment **L** ∈ ℝ^3^ due to the weight of the object when lifted off-center;
- *A clambed boundary condition at the Z* = 0 end of the trunk;
- *A free-end boundary condition at the Z* = *L* end of the trunk.

With these loading assumptions, the general form of eq. (11) and the kinematics in eq. (1) yield the following boundary value problem

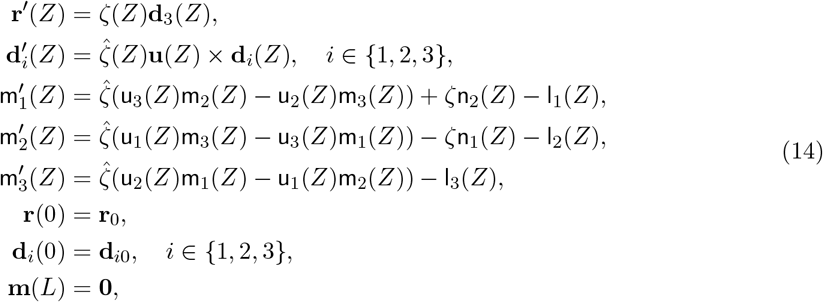

where {**d**_*i*_} form the director basis of the deformed configuration 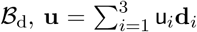 is the vector of curvatures of 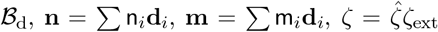 is the total extension of the rod in the deformed configuration, ζ_ext_ is the centerline extension due to the external loading relative to the activated configuration, **r**_0_ is the clamping position, the basis {**d**_*i*0_} defines the clamping direction, and

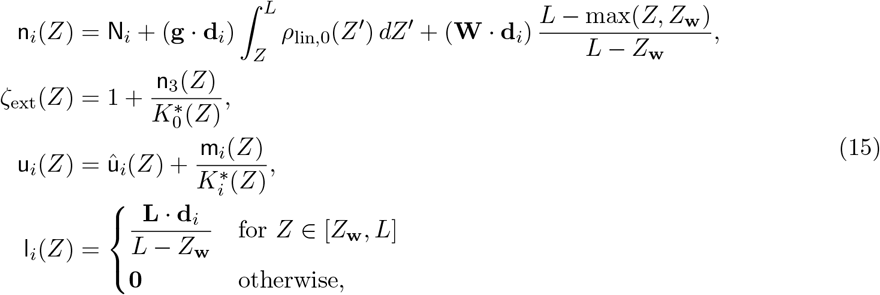

for *i* ∈ {1, 2, 3}, where 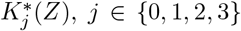,are the new stiffness coefficients in the activated configuration, ρ_lin,0_(*Z*) is the linear density along **e**_*Z*_ of the trunk in the initial configuration, and **g** = g**e**_*Z*_ is the gravitational acceleration. We assume that the contributions of body couples generated by the distributed load **w**(*Z*) are negligible due to the slenderness of the structure. Note that, to ensure consistency of centerline extensions across the initial, activated, and deformed configurations, we make the substitution 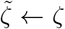 for the **r**^**′**^ differential equation and 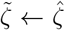 for the **d**^**′**^ differential equation in eq. (1) to express the kinematics in the boundary value problem in eq.(14).

We solve the differential system in eqs. (14) and (15) using the ‘colnew’ collocation method [60], which we found to outperform the shooting method equipped with various initial value problem solvers.

## 3. The elephant trunk representation

### Muscular subdomains

We can approximate the architecture of the elephant trunk and the associated fibrillar activation functions using a three-variable piecewise constant construction in *R*, Θ, and *Z*. In particular, we distinguish five muscle groups based on their fiber architecture and location in the cross section: dorsal longitudinal, outer ventral oblique, inner ventral oblique, dorsal radial, and ventral radial [38, 4]. We omit the transverse muscle architecture, since its mechanical contributions are equivalent to those of the radial muscle group and it is not straightforward to represent in a cylindrical basis. To represent accurately the trunk’s musculature while preserving model simplicity, we split the structure lengthwise into *N* = 3 piecewise segments ***Z*** ∈ [*Z*_1,*i*_, *Z*_2,*i*_], *i* ∈ {1, 2, 3}, and into symmetric right and left portions denoted by superscripts R and L, respectively. As a result, the following cylindrical regions define the 30 muscular subdomains that comprise our generalized representation of the elephant trunk:

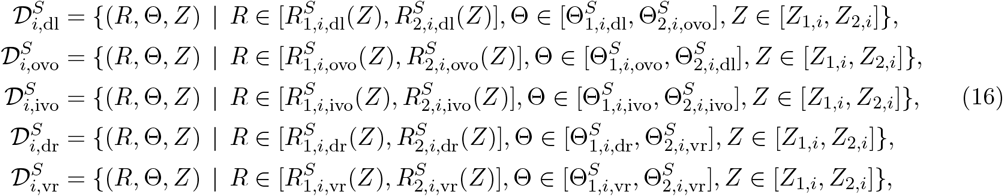

where *i* ∈ {1, 2, 3}, S ∈ {R, L}, ‘dl’ is the dorsal longitudinal group, ‘ovo’ and ‘ivo’ are the outer and inner ventral oblique groups, and ‘dr’ and ‘vr’ are the dorsal and ventral radial groups, respectively. That is, region 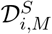 corresponds to the muscle group *M* ∈ {dl, ovo, ivo, dr, vr} on side *S* in the *i*-th piecewise *Z*-segment. The regions in eq. (16) assume that the inner and outer radii 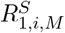 and 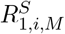 do not vary in Θ, so every cross section with a normal **e**_*Z*_ of a region 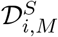 is an annular sector. To account for all other tissue surrounding the muscles of the trunk, each muscular subdomain 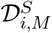 is a subset of the overall trunk region

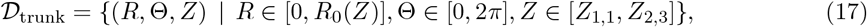

where *R*_0_(*Z*) is the outer radius of the trunk.

Based on the elephant trunk’s muscle architecture, we can assume that the left and right sides are symmetric along the axis going through the center of the dorsal longitudinal muscle region and the center of the cross-section. Under this assumption,

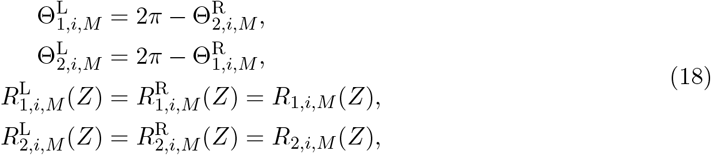

where *i* ∈ {1, 2, 3}, *Z* ∈ [*Z*_1,*i*_, *Z*_2,*i*_], and *M* ∈ {dl, ovo, ivo, dr, vr} indicates the muscle architecture and location. As such, we obtain a complete definition of the muscular subdomain geometry by prescribing 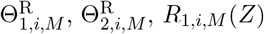 and *R*_2,*i*,*M*_ (*Z*).

### Fiber architecture

Following the definitions of the 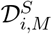 subdomains and assuming symmetry, we can fully describe the fiber architecture in the pre-tapered space 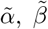as the following piecewise functions of **P** = (*R*, Θ, *Z*):

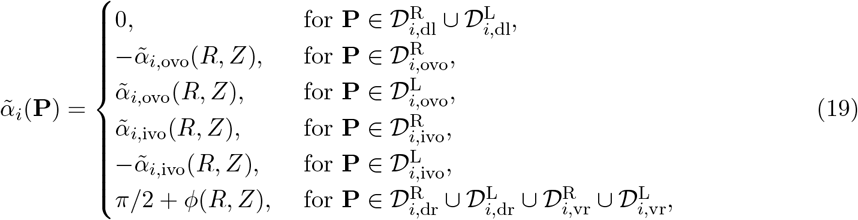

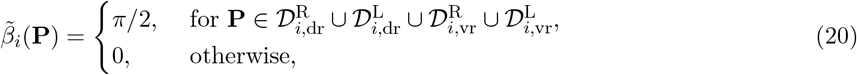

for *i* ∈ {1, 2, 3}, *Z* ∈ [*Z*_1,*i*_, *Z*_2,*i*_]. We emphasize that, in contrast to 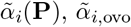,and 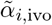are only functions of *R* and *Z*, and not Θ. Within a given oblique muscle region 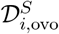 or 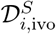 they obey the following geometric property that governs the helical angle variation in *R*:

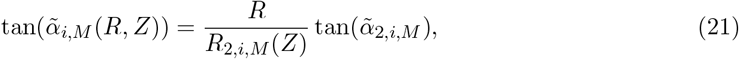

where *i* ∈ {1, 2, 3}, *Z* ∈ [*Z*_1,*i*_, *Z*_2,*i*_], *M* ∈ {ovo, ivo}, and 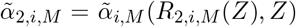 is a constant angle on the outer radius *R*_2,*i*,*M*_ (*Z*) of 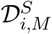 that fully defines a given helical architecture. Using eq. (21), we can then express the fiber architecture in eq. (19) in terms of the constant outer-helical angle quantities 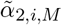.

Further, in the elephant trunk, the outer ventral oblique muscles are left-handed helical on the right side and right-handed helical on the left side. On the other hand, the inner ventral oblique muscles are right-handed helical on the right side and left-handed helical on the left side [38]. Therefore, we chose the negative signs in eq. (19) such that both 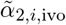 and 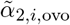 are positive. The tapering of the non-tapered fiber field given by 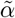 and 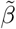 occurs through eq. (10) and the following rule for the inclination of the tapered field

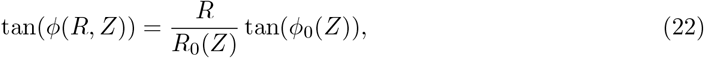

where *ϕ*_0_(*Z*) is the tapering angle on the outer surface of the trunk.

### Fiber activation functions

To prescribe the activation in all muscle groups, we construct a piecewise structure for the fibrillar activation function *ĝ*(*R*, Θ, *Z*) that is similar to eqs. (19) and (20). We assume that, within each muscular subdomain, the activation does not depend on *R*, which gives the general form

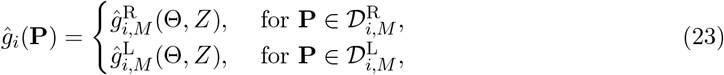

for *M* ∈ {dl, ovo, ivo, dr, vr}, *i* ∈ {1, 2, 3}, and *Z* ∈ [*Z*_1,*i*_, *Z*_2,*i*_]. Importantly, the symmetry assumption concerns only the muscular subdomain geometry and fiber architectures−the activation is generally different in the right and left sides of the trunk.

### Activated extension and curvatures

Based on the split of the integration regions into 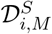 and the general form of activation formulas in eq. (6) for the disk solution, the *H*_0_, *H*_1_, *H*_2_, and *H*_3_ numerator terms of the activated extension and curvatures become piecewise functions in the three *Z*-segments such that

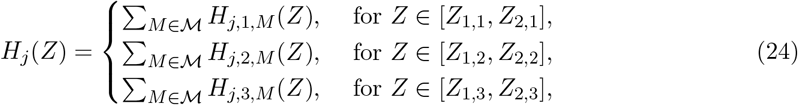

where *j* ∈ {0, 1, 2, 3}, ℳ = {dl, ovo, ivo, dr, vr}. We give all pertinent functional forms for eq. (24)

### Uniform activation and linear tabering

Assuming that the activation is uniform in each muscular subdomain, i.e., 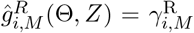 and 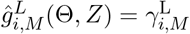 , the activation terms, provided in eq. (A.3) for general *ĝ*, simplify to

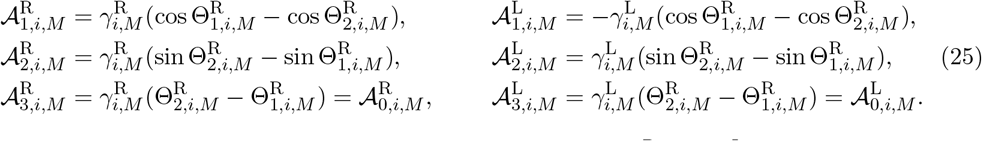

We note that, in the case of symmetric muscular activations 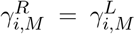 for given *i* and *M*, the corresponding activation terms obey the relations 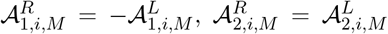 , and 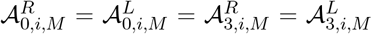 . In the case of symmetric 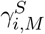 , these relations are due to the symmetries in the general form of eq. (A.3), which, in turn, stem from the symmetries assumed in eq. (18). We further assume that the tapering profile of the trunk representation is linear, which is a sufficiently good approximation of the geometry of a real trunk. In particular, we set the tapering angle evolution on the outer surface of the trunk to be constant, i.e., *ϕ*_0_(*Z*) = *ϕ*_0_. Then, the outer radius *R*_0_(*Z*) of the 𝒟_trunk_ region obeys the linear relationship

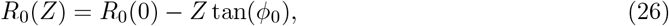

and the internal tapering of the fiber architectures in all muscular subdomains follows

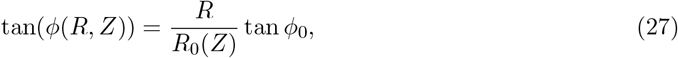

based on eq. (22). This form of *ϕ*(*R, Z*) enters the c_*ϕ*_ function needed to compute the *δ* functions in eq. (A.4). Further, for a linear tapering profile, we can readily express the linear density integral that enters eq. (15) as

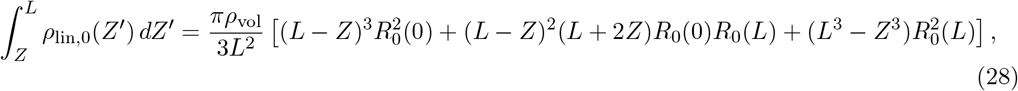

where ρ_vol_ is the volumetric density of the trunk material, assuming that ρ_vol_ is uniform throughout the trunk. We also make the assumption that the muscle fibers in a subdomain 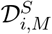 initiate at *Z*_1,*i*_ and terminate only at *Z*_2,*i*_, with no fibers terminating in the interior *Z*_1,*i*_ <*Z* < *Z*_2,*i*_ of each segment. With a non-zero tapering angle field *ϕ* (*R, Z*), this assumption imposes a condition on the tapered geometry of the muscular subdomains. Specifically, given the linearly tapered fiber field in eq. (27), we prescribe a compatibility condition

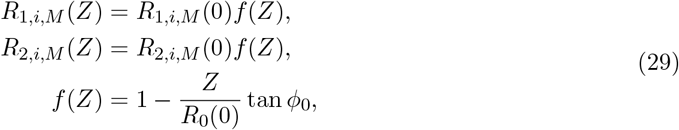

for *i* ∈ {1, 2, 3} and *M* ∈ ℳ, which ensures that the tangents of the inner surface *R* = *R*_1,*i*,*M*_ (*Z*) and the outer surface *R* = *R*_2,*i*,*M*_ (*Z*) of a muscular subdomain 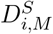 are the same as the fiber directions on those inner and outer surfaces, respectively, for all *Z* ∈ [*Z*_1,*i*_, *Z*_2,*i*_].

With the condition (29), we can fully define the trunk representation from 5+2*N* +(4·5)*N* +2*N* parameters. The five global parameters are: the Young’s modulus E; the Poisson’s ratio ν; the volumetric density ρ_vol_; the outer-surface tapering angle *ϕ*_0_; and the proximal trunk radius *R*_0_(0). The 2*N* parameters *Z*_1,*i*_ and *Z*_2,*i*_ define the bounds of the trunk segments. For each muscular subdomain we must specify the proximal radii *R*_1,*i*,*M*_ (0) and *R*_2,*i*,*M*_ (0), and the muscular subdomain angles 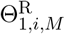 and 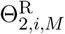 . For the oblique muscle groups, we need to define 2*N* helical angles 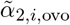 and 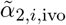.

### Image-informed trunk construction

We choose the values of the parameters that define the trunk representation based on the magnetic resonance images, dissection, and analysis of the trunk of an Asian elephant reported in [38]. First, the length of the imaged trunk is *L* = 1.75 m, and the ratio of the proximal outer radius to the total length is *R*_0_(0)/*L* ≈ 0.1185. We estimate the ratio of the proximal outer radius to the distal outer radius as *R*_0_(0)/*R*_0_(*L*) ≈ 3.46. Assuming a linear tapering profile, the outer-surface tapering angle then becomes *ϕ*_0_ = 4.81° based on *R*_0_(0), *R*_0_(*L*), and *L*.

Second, we choose *N* = 3 for the number of *Z*-segments, which results in a total of 30 independent muscular subdomains. We consider three segments to be sufficient to resolve, in an average sense, the otherwise highly complex muscular architecture and non-uniform activations of the real elephant trunk. Although choosing a much larger number of segments, such as *N* ∽ 10^2^, would result in an accurate correspondence with the real anatomy, it would significantly reduce the interpretability of muscular activation distributions, e.g., due to noisy activations extracted from the inverse problem. Further, a large *N* would increase the computational cost of inverse problem solutions, as the dimensionality of the problem is directly proportional to *N*.

We choose the segment coordinates as *Z*_1,1_ = 0, *Z*_1,2_ = *Z*_2,1_ = *L*/2, *Z*_1,3_ = *Z*_2,2_ = 0.85*L, Z*_2,3_ = *L*. The segment boundaries at *Z*_1,1_, *Z*_1,2_, and *Z*_1,3_ correspond to the three primary magnetic resonance images segmented along the length of the trunk in [38]. At each segment boundary, we redefine the muscular subdomain geometry through 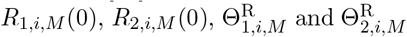 according to the three images. Based on the image that dictates the anatomy of the third segment, the inner ventral oblique muscle group is not present in that segment, which reduces the total number of muscular subdomains in the model to 28 and the total number of parameters to 72. Even though the third segment of the imaged trunk is very short, we elongated it to amount to 15% of the total length to emphasize the local effect of the non-existence of the inner ventral oblique muscles and to visualize the muscular contributions of the trunk’s tip in the simulations. We show the resulting 28-muscle representation of the trunk in fig. 3 and report the extracted geometry parameters in Table 1.

**Table 1:** Geometry parameters for the 28 muscular subdomains in three segments based on magnetic resonance images [38].

| | | segment $i = 1$ | segment $i = 2$ | segment $i = 3$ |
| --- | --- | --- | --- | --- |
| dorsal<br>longitudinal | $R_{1,i,dl}(0)/L$ | 0.0869 | 0.1016 | 0.0901 |
| | $R_{2,i,dl}(0)/L$ | 0.1185 | 0.1185 | 0.1185 |
| | $\Theta_{1,i,dl}^R$ | 0.0000 | 0.0000 | 0.0000 |
| | $\Theta_{2,i,dl}^R$ | 1.7279 | 2.0944 | 0.7854 |
| ----- |  | ----- | ----- | ----- |
| outer<br>ventral<br>oblique | $R_{1,i,ovo}(0)/L$ | 0.1027 | 0.1016 | 0.1024 |
| | $R_{2,i,ovo}(0)/L$ | 0.1185 | 0.1185 | 0.1185 |
| | $\Theta_{1,i,ovo}^R$ | 1.7279 | 2.0944 | 1.9635 |
| | $\Theta_{2,i,ovo}^R$ | 2.9845 | 2.9217 | 3.1416 |
| ----- |  | ----- | ----- | ----- |
| inner<br>ventral<br>oblique | $R_{1,i,ivo}(0)/L$ | 0.0869 | 0.0596 | — |
| | $R_{2,i,ivo}(0)/L$ | 0.1027 | 0.1016 | — |
| | $\Theta_{1,i,ivo}^R$ | 1.7279 | 1.0472 | — |
| | $\Theta_{2,i,ivo}^R$ | 3.1416 | 3.1416 | — |
| ----- |  | ----- | ----- | ----- |
| dorsal<br>radial | $R_{1,i,dr}(0)/L$ | 0.0580 | 0.0680 | 0.0579 |
| | $R_{2,i,dr}(0)/L$ | 0.0869 | 0.1016 | 0.0876 |
| | $\Theta_{1,i,dr}^R$ | 0.0000 | 0.0000 | 0.0000 |
| | $\Theta_{2,i,dr}^R$ | 1.5708 | 1.0472 | 0.7854 |
| ----- |  | ----- | ----- | ----- |
| ventral<br>radial | $R_{1,i,vr}(0)/L$ | 0.0470 | 0.0281 | 0.0318 |
| | $R_{2,i,vr}(0)/L$ | 0.0761 | 0.0596 | 0.0766 |
| | $\Theta_{1,i,vr}^R$ | 1.5708 | 1.0472 | 1.5708 |
| | $\Theta_{2,i,vr}^R$ | 3.1416 | 3.1416 | 3.1416 |

**Figure 3:**
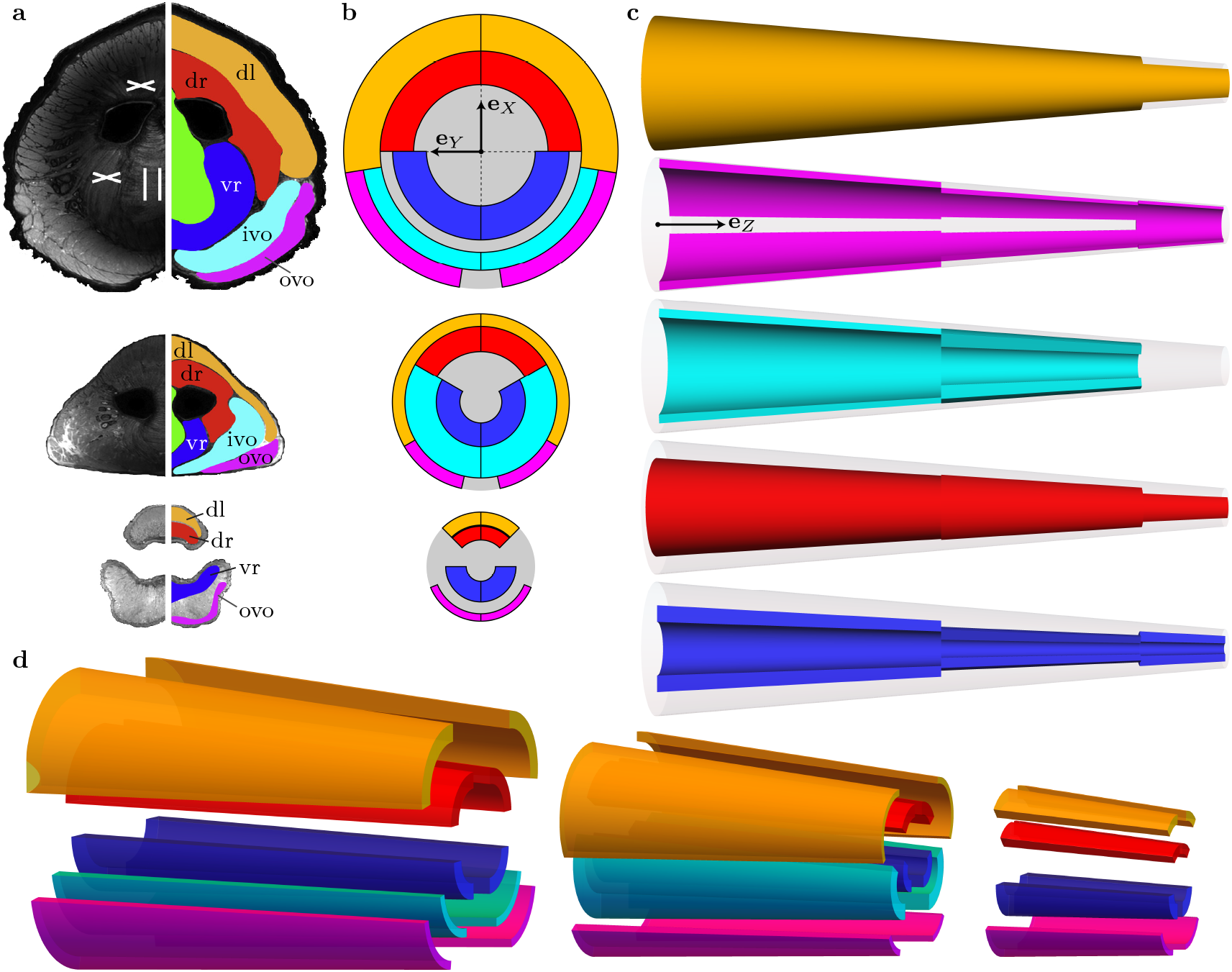
Trunk model geometry constructed from cross-sectional magnetic resonance images of the trunk of an Asian elephant. (a) Magnetic resonance images of transversal sections of the trunk (left column), and the corresponding segmented muscle groups (right column) at three points along the length of the trunk. The magnetic resonance images and the muscle segmentation are adapted from [38]. (b) Cross sections of the assumed muscular subdomains 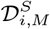 based on the images in (a). The three cross sections correspond to the three images in (a) and to the starting points *Z*_1,1_ (top), *Z*_1,2_ (middle), and *Z*_1,3_ (bottom) of the three segments in the model. We omit the transverse muscle group shown in green in (a). (c) Dorsal visualizations of the five muscle groups, from top to bottom, in all three segments of the trunk model. (d) Exploded view of all 28 muscular subdomains used in the trunk model. We use the same color coding of all muscles across the subfigures (a)-(d).

Finally, we used the upper bound of the 20°–30° range for the fiber angles provided in [38], such that the unsigned helical angles are 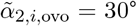 for *i* ∈ {1, 2, 3}, and 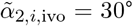 for *i* ∈ {1, 2}. We assume a homogeneous incompressible material with *ν* = 0.5, a Young’s modulus of *E* = 1 MPa [43], and a volumetric density of *ρ*_vol_ ≈ 1059.7 kg/m^3^ [61].

### Incombressibility effects

The elephant trunk is a muscular hydrostat, and its incompressibility is critical to enable various biomechanically desirable phenomena [1, 55]. For instance, contraction of non-longitudinal muscle groups can directly reduce the cross-sectional area which, in turn, results in axial extension due to incompressibility. Combined with other mechanical effects, the elephant trunk can exploit this mechanism to reach extensional strains larger than 30% [38]. As another example, the asymmetric contraction of radial muscles can induce localized reduction in cross-sectional area leading to an extension mismatch in the cross section which, in turn, causes bending.

Equations (5)-(7) and (24)-(A.4) already incorporate some of the hydrostatic effects, as the activation terms *H*_*j,i,M*_ depend on the Poisson’s ratio *ν*. For instance, eqs. (24)-(A.4) capture the bending deformation mode induced by radial muscles when *ν* = 0.5. However, those expressions alone do not consider the change in the geometry of the activated configuration caused by the hydrostatic effects. To incorporate the change in the outer radius profile due to incompressibility, we assume that the muscular contractions throughout *Z* ∈ [0, *L*] have an integral effect on the cross-sectional size of the trunk at all *Z*. Specifically, we scale the outer radius profile *R*_0_(*Z*) by a prefactor dependent on the muscular activation, so that the volumes of the activated and initial configurations are the same. Under the assumption of linear tapering, scaling *R*_0_(*Z*) by a constant prefactor implies that the activated configuration remains a conical frustum, which effectively averages the localized effects of the otherwise variable extension 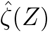.To ensure that the volumes of the conical frustums are the same before and after activation, we employ the following scaling to compute the outer radius 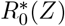 in the activated configuration

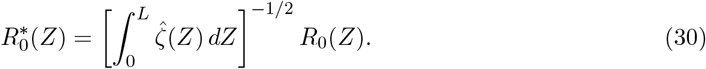

We use this updated radius profile 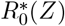 to compute the stiffness coefficients 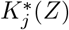 of the activated configuration, which enter eq. (15). For visualization of the final configuration with external loading, we use eq. (30) with ζ(*Z*) in place of 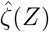 to compute the outer radius profile of **ℬ**_d_.

We emphasize that the computation of the activated curvature and extension according to eq. (5) is purely analytical, given our stated assumptions. That is, obtaining û(*Z*) and 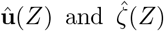 does not require any iterative procedures, and we can consider computing these functions to be a nearly instantaneous operation. This analytical aspect of the computation of **ℬ** enables real-time prediction of the deformations due to muscular activation in the absence of external loading. We can also achieve real-time computation of the externally loaded configuration **ℬ**_d_, which requires a numerical solution to eqs. (14) and (15), by optimizing the computational structure of the model and implementing precomputation measures wherever possible.

First, we observe that the functions *δ*_j,*i*,*A*_ and *K*_j_ do not depend on the muscular activations 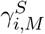, for all *i* ∈ {1, 2, 3}, *j* ∈ {0, 1, 2, 3}, A ∈ {lo, he, ra}, *M* ∈ ℳ, and S ∈ {R, L}. Thus, we precompute all *K*_j_ and *δ*_j,*i*,*A*_, evaluated at arguments given in eq. (A.2), with uniform discretizations over the intervals *Z* ∈ [*Z*_1,*i*_, *Z*_2,*i*_] consisting of n points, such that

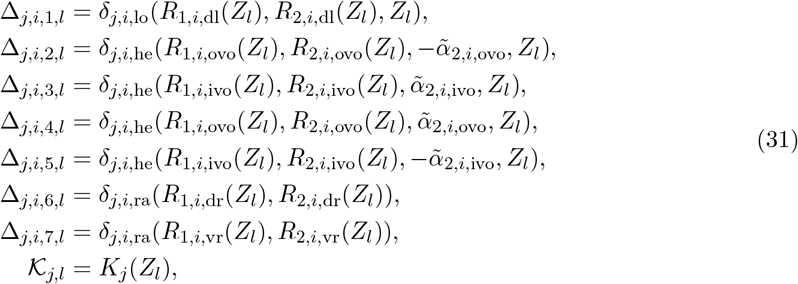

for *Z*_l_ = *Z*_1,*i*_ + *l*(*Z*_2,*i*_ − *Z*_1,*i*_)/(n — 1), *l* ∈ {0,…, n − 1}. We note that, due to the symmetry conditions in eq. (18) and the properties of the assumed muscle architectures, eq. (31) needs to define only seven architectural classes of discretized Δ quantities required for precomputation.

Second, we can decompose the fractional terms *H*_j_/*K*_j_ in eq. (5) as

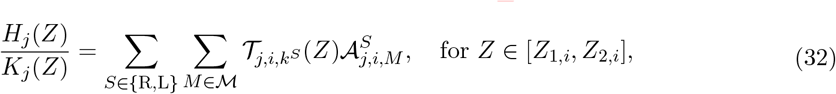

where 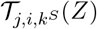 are cubic interpolating functions over *Z* ∈ [*Z*_1,*i*_, *Z*_2,*i*_] with values

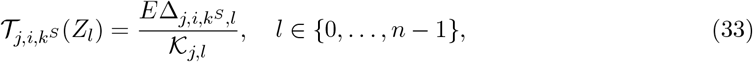

at the control points, and *k*^R^ = *k*^L^ = 1 for *M* = dl, *k*^R^ = 2 and *k*^L^ = 4 for *M* = ovo, *k*^R^ = 3 and *k*^L^ = 5 for *M* = ivo, *k*^R^ = *k*^L^ = 6 for *M* = dr, *k*^R^ = *k*^L^ = 7 for *M* = vr. We can then use the interpolating functions 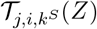 in their precomputed form regardless of the applied fibrillar activation. Therefore, after the precomputation phase, we only need to obtain the trivial activation terms 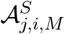 in eq. (25) for a given set of muscular contractions before proceeding with the numerical solution of the boundary value problem. As a result, the computation time for the trunk deformation with external loading and arbitrary muscular activation is only on the order of 0.1 seconds on a standard desktop computer.

### Motion of the trunk’s broximal base

While an elephant relies primarily on the deformation of its trunk due to the internal trunk muscles, many motion tasks involve movement of the elephant’s head which, in turn, moves and rotates the proximal base of the trunk. We model the motion of the base of the trunk as an evolution of the boundary conditions **r**_0_ and **d**_*i*0_, *i* ∈ {1, 2, 3}, in eq. (14). In particular, we assume that the motion of the trunk base occurs on a sphere of radius *r*_S_ ≈ *L*/4 based on the approximate anatomical distance from the proximal end of the trunk to its center of rotation located approximately at the cranial end of the elephant’s neck. To provide a reference for subsequent transformations that govern the motion and rotation of the trunk’s base, we define the default location **r**_𝒟_ = **0** of the base and a constant center **S** of the sphere. In particular, the center of the sphere is always at

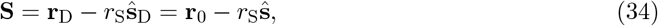

where **ŝ**_D_ is a unit vector pointing from the center of the sphere to the default location of the base, while **ŝ** is a unit vector from **S** to **r**_0_ at a given time point. We express the default vector **ŝ**_D_ through the transformation

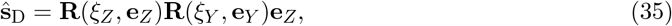

where **R**(ξ, **e**) denotes a rotation transformation by an angle ξ about an axis **e**, and the constant angles ξ_*Z*_, ξ_*Y*_ effectively define **S** relative to the default location **r**_D_. Then, we can define the motion as **r**_0_ = **S** + *r*_S_**ŝ** with

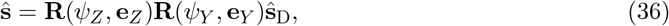

where *ψ*_*Z*_ and *ψ*_*Y*_ are the spherical angles controlled by the motion of the elephant’s head that describe the position of the trunk’s base on the sphere. Since the centerline tangent **d**_30_ at the base of the trunk is not necessarily normal to the sphere, we introduce an angle ξ_**d**_ that defines a set of default clamping directions **ŝ**_**d**, D,*i*_ for the directors **d**_*i*0_, i.e.,

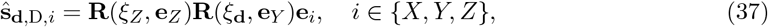

and the following dependence of the director basis at *Z* = 0 on the motion angles *ψ*_*Z*_ and *ψ*_*Y*_ :

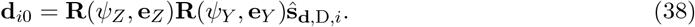

As a result, eq. (37) prescribes a constant angular offset between the trunk base directors **d**_*i*0_ and the sphere tangent **R**(*ψ*_*Z*_, **e**_*Z*_)**R**(*ψ*_*Y*_ , **e**_*Y*_ )**e**_*Z*_ at **r**_0_, for any *ψ*_*Z*_ and *ψ*_*Y*_ . This ensures the rigidity of the orientation of the trunk’s base relative to the elephant’s head. We inform the value of ξ_**d**_ by the musculoskeletal anatomy of the elephant’s head, which gives an approximation ξ_**d**_ ≈ 30°. We set the default position of the trunk’s base as ξ_*Z*_ = 0° and ξ_*Y*_ = 100°.

We note that applying another fixed offset rotation to **d**_*i*0_, in addition to **R**(ξ_**d**_, **e**_*Y*_ ), is not necessary to fully define the geometry of the system since the tangent direction of the trunk’s base lies in the sagittal symmetry plane of the elephant’s head. Further, given that we consider two rotation angles *ψ*_*Z*_ and *ψ*_*Y*_ in eqs. (36) and (38) rather than three, we effectively assume that the motion of the elephant’s head does not induce a twist of the director basis around the normal vector **ŝ**.

## 4. Biomechanical principles of the trunk representation

The analytical results in eqs. (5) and (A.1)-(A.4) as well as the physiological parameters in Table 1 provide insights into the fundamental biomechanical principles that govern the elephant trunk motion. Specifically, we can infer the isolated mechanical effect, e.g., the induced deformation modes and their directions, for each individual muscle group. In this section, we present a quantitative discovery of muscle group-specific effects by using our trunk model.

To investigate the effect of each individual muscle group on the trunk’s deformation, we analyze the contribution of a given group to the extension 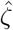,the curvatures û_1_ and û_2_, and the twist density û_3_. Since muscular activation is contractile, we assume that the uniform activations 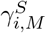 that enter eq. (25) are non-positive. Due to the additive property of the *H*_j,*i*,*M*_ functions in eq. (24), we quantify the contribution of a muscle group *M* to 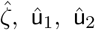 ,and û_3_ using the terms *H*_0,*i*,*M*_ /*K*_0_, *H*_1,*i*,*M*_ /*K*_1_, –*H*_2,*i*,*M*_ /*K*_2_, and *H*_3,*i*,*M*_ /*K*_3_, respectively. We observe that, given eq. (25) and the angular bounds 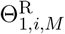 and 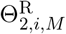 of the muscular subdomains in Table 1, the signs of the individual contributions of each subdomain to any given deformation mode are the same in all segments and for all *Z* within a given segment. Thus, we can discuss the directions of the mechanical effects of each muscle type for all three segments and for all *Z* ∈ [0, *L*] simultaneously.

Further, we introduce the torsion 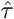 of the activated trunk configuration as

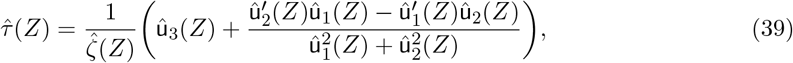

which characterizes the non-planarity of the trunk’s centerline [56]. The cases of 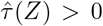 and 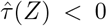 correspond, respectively, to the right-handed and left-handed helicity of the trunk’s activated shape at a given *Z*. On the other hand, the sign of û_3_ describes the direction of twist of the trunk around the tangent 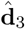.

It is important to note that while we can discuss the contributions of individual muscle groups to 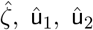 and û_3_ in an additive manner—e.g., conclude that the dorsal longitudinal muscles increase û_2_ upon contraction with other arbitrary muscular contractions present—the changes in torsion due to additional muscular activations are nonlinear and nontrivial due to the curvature-torsion coupling. For instance, the torsion can generally change due to activation of the dorsal longitudinal muscles, even though these muscles produce no torsion in the absence of other muscular contractions. Therefore, we consider the *isolated torsion* contributed by each muscle group, i.e., the torsion generated by a given muscle group with no other muscular activations in the trunk.

From our analysis, we directly obtain that the right and left dorsal longitudinal muscles always induce negative and positive bending around 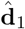, respectively, and only positive bending around 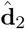. The longitudinal muscles also do not introduce any twist and they do not generate any torsion on their own. The outer ventral oblique muscles have the same effect on bending around 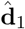, as the dorsal longitudinal group, but the opposite effect on bending around 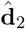, i.e., both the right and left outer ventral oblique muscle groups have negative contributions to û_2_. In addition, the right outer ventral oblique muscles induce positive twist and torsion, while the left muscles contribute negative twist and torsion. The signs of the contributions of the inner ventral oblique muscles to û_1_ and û_2_ are the same as for the outer ventral oblique group. The difference between the two groups lies in the direction of the twist and torsion—the inner ventral oblique muscles contribute twist and torsion of the opposite sign compared to the outer ventral oblique muscles. In contrast, the radial muscle groups do not contribute any twist and do not produce any torsion in isolation. However, while the longitudinal and oblique muscles cause overall contraction of the trunk’s centerline, the radial muscles extend the trunk due to the muscular-hydrostat effect. Further, both dorsal and ventral radial muscle groups contribute bending around 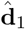 in the opposite direction compared to both the longitudinal and oblique muscles. For bending around 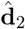, both right and left dorsal radial muscles have a negative contribution, while both right and left ventral radial muscles have a positive contribution to û_2_. We summarize all these findings in Table 2.

**Table 2:**
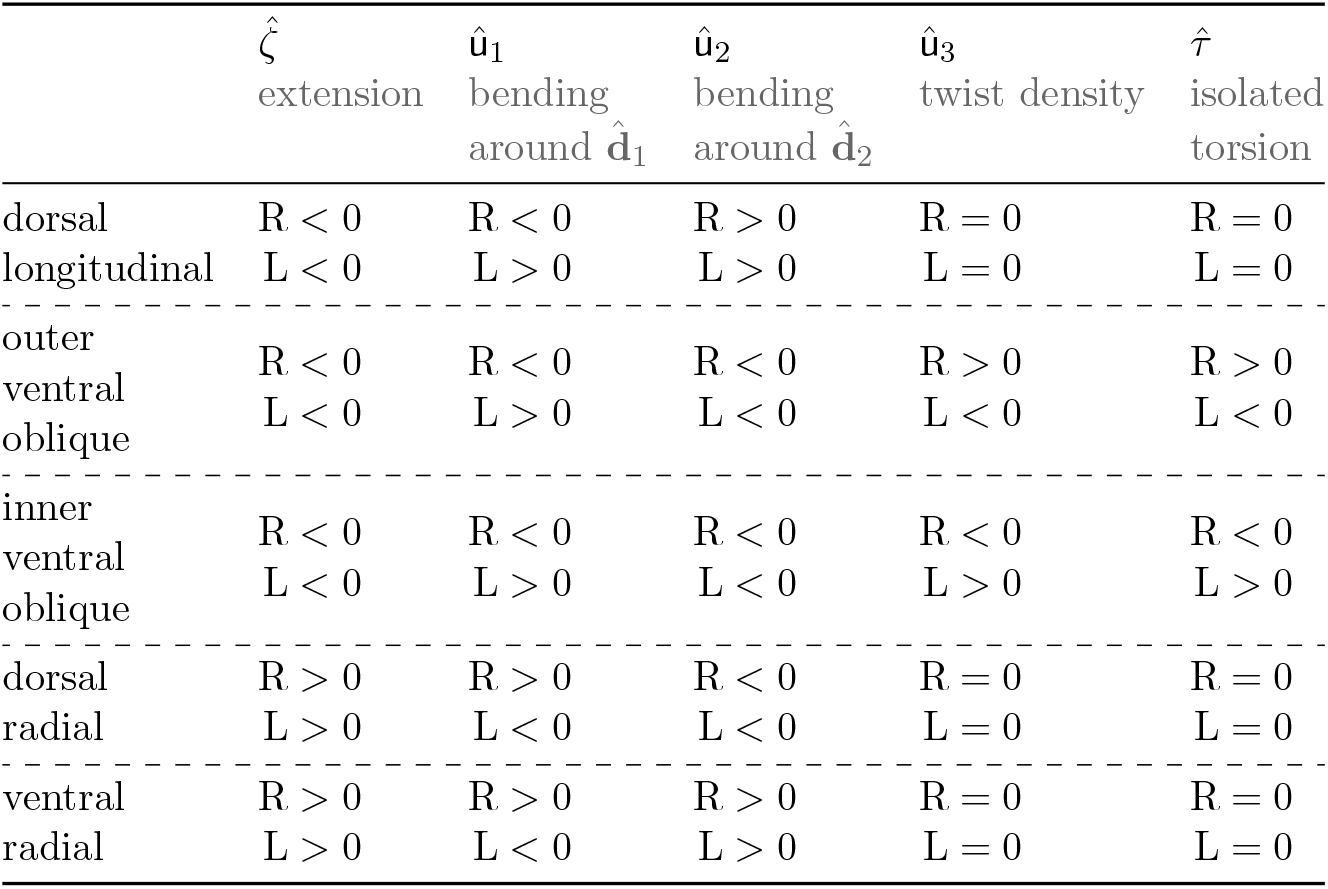
Contributions of individual muscle groups to the deformation of the elephant trunk. The map derives from the analytical elephant trunk model and assumes uniform fibrillar activation within a given muscular subdomain. The designations ‘R’ and ‘L’ correspond to the contributions of the right and left trunk muscle groups, respectively.

The derived properties of the muscular contributions are consistent with the intuition that the shortening of the dorsal section causes the trunk to bend around 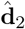, in the dorsal direction towards 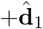, while shortening of the right and left sections induces rightward and leftward bending around 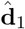, towards 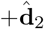 and 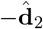, respectively. Shortening of the ventral section has the same effect on bending around 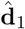 as that of the dorsal section; however, ventrally located muscles lie on the opposite side of the 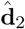 axis, which reverses their contributions to bending around 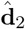. The directions of all these effects further reverse when we replace the shortening process with an elongation, which occurs in radial muscles that indeed demonstrate a reversed set of contribution signs. For oblique muscles, the signs of the mechanical effects also depend on their helical angles 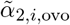 and 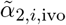. In particular, the signs of the effects on the twist and isolated torsion reverse when the sign of the helical angle changes; hence the opposite signs for the û_3_ and 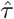 contributions between the outer and inner oblique muscles. Additionally, above a certain helical angle magnitude 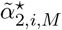 for a given muscular subdomain [54], the signs of the contributions to 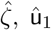 and û_2_ flip. The signs for the twist and isolated torsion contributions remain the same regardless of the helical angle magnitude.

We note that the critical helical angles 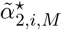 are conceptually the same as the special angle of 54.73° encountered in McKibben actuators [62]. However, in the model considered here, they are no longer a single constant since the tapering of the fiber fields and the helical angle variation in eq. (21) render them more involved functions of *ϕ*_0_ as well as *R*_1,*i*,*M*_ (*Z*) and *R*_2,*i*,*M*_ (*Z*). All oblique muscular subdomains in the particular trunk representation constructed here do not exceed their corresponding critical helical angle magnitudes, i.e., 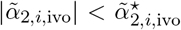 and |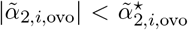 , for all *i* ∈ {1, 2, 3}.

The analysis so far considered only the signs of the contributions of the individual muscle groups. The complete representation of the elephant trunk, however, also provides an explicit ranking of the magnitudes of the contributions to 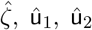 and û_3_ for a given muscular activation γ held constant across all groups. Further, as a consequence of assuming a constant tapering angle, we can apply to all *Z* in a given segment any conclusion regarding the contribution magnitudes derived at one particular *Z* in that segment. In Table 3, we report the ranking of the muscle groups within each segment sorted according to the magnitudes of their contributions |*H*_0,*i*,*M*_ /*K*_0_|, |*H*_1,*i*,*M*_ /*K*_1_|, |*H*_2,*i*,*M*_ /*K*_2_|, |*H*_3,*i*,*M*_ /*K*_3_| to 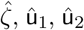 and û_3_, respectively. For a fixed range of muscular activations, the dorsal longitudinal muscles dominate the bending curvatures û_1_ and û_2_ in all segments, but they cannot provide any twist. The oblique muscles are the most versatile, as they contribute high-magnitude effects to both bending in û_1_ and û_2_ as well as twisting in û_3_.

**Table 3:** Ranking of the magnitudes of individual muscle group contributions to the deformation of the trunk, as predicted by the model. Within each *Z*-segment, we order the five muscle groups according to the magnitudes, from lowest to highest as indicated by the arrows, of their corresponding contributions to different deformation measures. For adequate comparison, all contributions result from the same muscular activation magnitude. The ‘(0)’ designation indicates that the given muscle groups do not contribute to that measure.

| | $\hat{\zeta}$<br>$\vert H_{0,i,M}/K_0\vert $ | $\hat{u}_1$<br>$\vert H_{1,i,M}/K_1\vert $ | $\hat{u}_2$<br>$\vert H_{2,i,M}/K_2\vert $ | $\hat{u}_3$<br>$\vert H_{3,i,M}/K_3\vert $ |
| --- | --- | --- | --- | --- |
| Segment $i = 1$ | ↑ dl | dl | dl | ovo |
|  | dr | ovo | ovo | ivo |
|  | ovo | ivo | ivo | vr, dr, dl (0) |
|  | vr | dr | dr |  |
|  | ivo | vr | vr |  |
| ----- |  |  |  |  |
| Segment $i = 2$ | ↑ ivo | dl | dl | ivo |
|  | dl | ivo | ivo | ovo |
|  | dr | ovo | dr | vr, dr, dl (0) |
|  | vr | dr | ovo |  |
|  | ovo | vr | vr |  |
| ----- |  |  |  |  |
| Segment $i = 3$ | ↑ dl | dl | dl | ovo |
|  | vr | ovo | ovo | vr, dr, dl (0) |
|  | ovo | vr | vr |  |
|  | dr | dr | dr |  |

The contributions to bending of both dorsal and ventral radial muscles are the smallest, which agrees with the intuition that the local elongation due to the muscular-hydrostat effect has a smaller impact on the bending curvature than the shortening due to direct contraction of longitudinal fibers. Further, the smaller contributions of the radial muscles to bending are also consistent with the fact that the radial muscle groups reside closer to the central axis of the trunk. In contrast, the longitudinal muscles are closer to the trunk’s outer surface, thus requiring smaller muscular activations to achieve the same extent of bending. Finally, between the three segments, the ranking of effect magnitudes for the extension 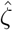 is the most variable primarily due to the different cross-sectional areas of each muscle type across the three segments.

## 5. Construction of trunk motions

### The inverse motion problem

Next we seek to identify the underlying muscular activations in the elephant trunk model during a set of given motion tasks. While previous sections tackled the forward problem of computing the deformed configuration for a given trunk design and muscular activations, finding the muscular activations that achieve a desired motion is an inverse problem. Previous work investigated various inverse problems in the control and design of active slender structures [63, 64], particularly how they become ill-posed when the number of independently activatable fibrillar regions is greater than three [65]. Importantly, the fibrillar activation solution to an inverse problem with four or more activation regions can generally become non-unique, meaning an infinite number of fibrillar activation arrangements can reach the same point in space.

With 28 muscular subdomains, our trunk representation is clearly a highly redundant system capable of achieving a given functional task, such as moving objects between two points, with an infinitely diverse variety of muscular activity. While the physiological constraints on the muscular contraction magnitudes limit this diversity, the trunk representation still provides a vast set of potential fiber contraction arrangements that yield the same functional result. In this work, we seek to identify a single solution of the inverse problem for a given motion task, while acknowledging that a large family of other solutions can be equally valid.

### Optimization method

In addition to the ill-posedness aspect, the complexity of eqs. (14) and (15) implies that a closed-form solution to the inverse problem is not possible. Therefore, we resort to an automatic optimization method to obtain one possible solution to the inverse problem.

In particular, we define a given motion task as a function 𝒫 that defines discrete geometrical properties at points *Z*_p_ ∈ [0, *L*] of the deformed configuration **ℬ**_d_ that we would like the trunk to achieve at a discrete time index *T* ∈ {1,…, *T*_max_},

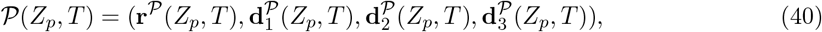

for *p* ∈ {1,…, *N*_p_}, where **r**^*𝒫*^ (*Z*_p_,*T* ) is the desired time evolution of the centerline point **r** at *Z*_p_, and 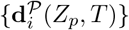 is the desired evolution of the director basis {**d**_*i*_} at *Z*_p_. We assume that every motion starts at a fixed state 𝒫(*Z*, 0) defined by zero fibrillar activation, boundary conditions with **ψ**_*Z*_ = **ψ**_*Y*_ = 0, and only the external loading due to the weight of the trunk. Further, we will use ‘○’ in place of any of the four elements in 𝒫(*Z*_p_,*T* ) to denote that the corresponding element of 𝒫(*Z*_p_,*T* ) can take on arbitrary values, and it is not relevant to the motion.

To identify the muscular activation that leads to a quasi-static motion 𝒫, we solve the following sequence of optimization problems for *T* ∈ {1,…, *T*_max_}:

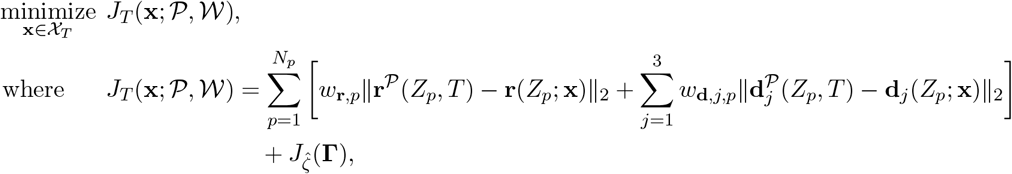

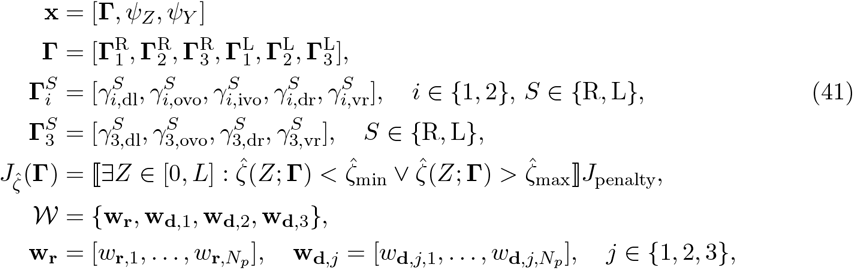

where **x** ∈ ℝ^30^ is a vector with respect to which we minimize the objective function *J*_*T*_ at time *T*, χ_*T*_ is the feasible set for **x** at time *T, w*_**r**,*p*_ is the weight for the Euclidean norm deviation in **r** at *Z*_*p*_, *w*_**d**,*j*,*p*_ is the weight for the Euclidean norm deviation in **d**_*j*_ at *Z*_*p*_, {**r**(*Z*; **x**), **d**_1_(*Z*; **x**), **d**_2_(*Z*; **x**), **d**_3_(*Z*; **x**)} is the solution to the boundary value problem in eqs. (14) and (15) computed for a given activation ***Γ*** and trunk base rotation {**ψ**_*Z*_, **ψ**_*Y*_}, 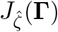 is a penalty term added whenever 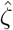 violates the ζ_min_ *<* ζ *<* ζ_max_ condition at any *Z* ∈ [0, *L*] for a given activation ***Γ***, ⟦*P*⟧ is the Iverson bracket returning 1 if *P* is true and 0 if *P* is false, and *J*_penalty_ > 0 dictates the magnitude of the additional penalty term.

In other words, the objective function in eq. (41) measures a weighted *L*_2_-norm deviation between the deformed configuration ℬ_*d*_ and the desired properties of that configuration at a time *T*. Finding **x** ∈ χ_*T*_ such that *J*_*T*_ ≈ 0 corresponds to identifying the muscular activations and the trunk base rotation that match the desired properties of the deformed shape. Including the penalty term 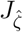 ensures that the computed muscular activations do not produce a non-physiological axial strain in any of the three *Z*-segments. In our simulations, we set 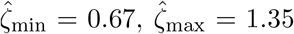 based maximum contractile and extensile strains observed physiologically in elephant trunks [43, 38], with a penalty factor of *J*_penalty_ = 10^3^.

We solve the optimization problem at each time *T* using the Julia Optimization.jl package [66] by using either (1) the NOMAD blackbox optimization scheme [67] from the NOMAD.jl package [68] for global optimization, or (2) the Sbplx scheme [69, 70] from the NLopt library [70] for local optimization in a close neighborhood of the initial guess. We prescribe the feasible set χ_*T*_ as a hyperrectangle defined by the intervals [*x*_*i*,min_(*T*), *x*_*i*,max_(*T*)] for each of the 30 components of **x**. While we explicitly choose the values of *x*_*i*,min_(0) and *x*_*i*,max_(0) for the first time point, the feasible sets χ_*T*_ at all subsequent time points *T* generally depend on the optima found at the previous time points *T* − 1 or optima computed without external loading. For all optimization problems in this work, we restrict the muscular activations to be contractile only, such that *x*_*i*_ ≤ 0, *i* ∈ {1,…, 28}, and to not exceed a prescribed maximum activation magnitude |*γ*|_max_, i.e., |*x*_*i*_| ≤ |*γ*|_max_, *i* ∈ {1,…, 28}. Defining sufficiently constrained feasible sets is critical to achieve computational feasibility in solving the optimization problem. Refer to Appendix B and Appendix C for a more detailed description of how we construct the feasible sets, choose the initial guesses, and select the optimization schemes to compute the trunk motions.

Since 𝒫 is discrete over the time indices *T*, we interpolate the muscular activations and trunk base rotations between subsequent indices *T* with a continuous time parameter *t* to generate a continuous motion of the trunk. In particular, we define a time interval *t* ∈ [*t*_*i*_, *t*_*i*+1_] for any pair of discrete time indices (*i, i* + 1), *i* ∈ {0,…, *T*_max_ − 1}. We then compute the continuous motion between *T* = *i* and *T* = *i* + 1 using an interpolation **x**(*t*) = ℐ_*i*_(*t*; *t*_*i*_, *t*_*i*+1_, **x**^*\**^(*i*), **x**^*\**^(*i* + 1)) over *t* ∈ [*t*_*i*_, *t*_*i*+1_] with values **x**^*\**^(*i*) at *t*_*i*_ and **x**^*\**^(*i* + 1) at *t*_*i*+1_, where **x**^*\**^(*i*) and **x**^*\**^(*i* + 1) are the outputs from the minimization of *J*_*i*_ and *J*_*i*+1_, respectively. We specify the particular form of ℐ_*i*_ for each analyzed motion independently, depending on the qualitative characteristics of the motion.

Critically, we note that the constraints on the activations **Γ** and the angles **ψ**_Z_, **ψ**_Y_ , as well as the specificity of a given motion definition 𝒫 all serve to isolate a desired set of configuration shapes in an otherwise vast family of possible trunk motions that achieve 𝒫. For any particular deformation at a given time point within a motion, there are generally infinitely many other activation sets that achieve the exact same deformation due to redundancy. Apart from the activation-level redundancy, there are generally infinitely many sets of deformations {**r**(*Z*), **d**_1_(*Z*), **d**_2_(*Z*), **d**_3_(*Z*)} that exactly fulfill a given motion definition 𝒫. As such, to guide the optimization process, we take special care in designing both the definitions 𝒫 and the feasible sets, so that the motions computed through optimization bear physiological resemblance. Upon choosing 𝒫 and the feasible sets, we seek to identify, at each *T*, the most desirable **x**^*\**^(*T*) that is not only a minimum of *J*_T_ , but rather a global minimum that also results in a sufficiently small value *J*_T_ (**x**^*\**^(*T*); 𝒫, 𝒲). In the case that even the *global* minimum produces a *J*_T_ that indicates insufficient matching of the desired configuration, we revise the feasible set to allow other minima to emerge.

## 6. Analysis of three motion tasks

### Picking and eating a fruit

The first elephant trunk motion that we simulate involves picking a fruit from a tree branch and eating it. We place the fruit at a high position relative to the distal end of the trunk to explore the muscular contractions needed for the trunk to overcome its own weight in upward-bending configurations. The fruit picking motion consists of three phases: (1) reaching the fruit location with the tip of the trunk, (2) increasing the pulling force on the fruit until a prescribed threshold tension force, (3) picking the fruit from the branch and moving it to the elephant’s mouth.

We define the first phase as

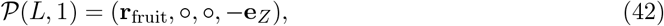

where **r**_fruit_ is the location of the fruit, with the loading conditions consisting of only gravitational forces acting on the trunk, i.e., **N** = **W** = **L** = **0**. Equation (42) guides the trunk to reach the point **r**_fruit_ with its distal end at *Z* = *L* while enforcing the centerline tangent **d**_3_ at the distal end to point upwards. The functions 𝒫 for the second phase,

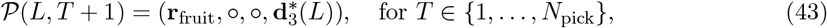

are almost the same as in eq. (42), with the exception that the desired centerline tangent is the optimum 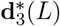 established by minimizing *J*_1_. The motion in the second phase consists of multiple states 𝒫(*L, T* + 1), *T* ∈ {1,…, *N*_pick_}, where *N*_pick_ is the number of intermediate states for the increasing tension exerted on the fruit during the picking process. In particular, we set 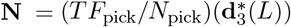, where *F*_pick_ is the threshold tension that leads to separation of the fruit from the branch. We note that, during the picking process, the weight of the fruit does not cause a force on the trunk, since the fruit is still hanging from the branch. Further, in the picking phase, we also have **W** = **L** = **0**. In the final phase, the fruit detaches from the branch, and the motion ends according to

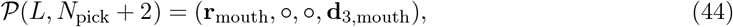

where **r**_mouth_ is the approximate location of the elephant’s mouth, and **d**_3,mouth_ is a desired orientation of the distal end of the trunk at the mouth location. The weight of the fruit itself causes a force **N** = *m*_fruit_*g*e_Z_ acting on the endpoint *Z* = *L* of the trunk, where *m*_fruit_ is the mass of the fruit. The tension force from the picking phase is no longer present in the final phase.

We define the particular fruit-picking motion evaluated here with the following parameter choices: **r**_fruit_ = [0.7 m, 0.3 m, −0.2 m], *F*_pick_ = 100 N, **r**_mouth_ = [−0.2 m, 0 m, 0.16 m], **d**_3,mouth_ = **R**(−3*π*/4, **e**_Y_ )**e**_Z_, *t*_0_ = 0, *t*_*i*_ = (3/2) + (*i* − 1)(2/3)*/N*_pick_ for *i* ∈ {1 .. ., *N*_pick_ + 1}, 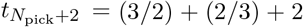, *N*_pick_ = 20, and *m*_fruit_ = 8 kg. As a result, the durations of the three motion phases are approximately 1.5, 0.67, and 2.0, respectively. We use cosine interpolation for both ℐ_0_ and 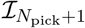, and no interpolation for ℐ_*i*_, *i* ∈ {1,…, *N*_pick_}, since the number of explicit intermediate points *N*_pick_ = 20 is sufficient to approximate a continuous motion between *T* = 2 and *T* = *N*_pick_ +1. We use an exaggerated mass *m*_fruit_ = 8 kg for the fruit to emphasize the effects of its mass on the motion and demonstrate the ability of the model to handle external loads. Throughout the three phases, 𝒫 acts only on the endpoint *Z* = *L*, so *N*_p_ = 1 in the definition of the optimization problem. We set |γ|_max_ = 3.0 and assign the weights **w**_**r**_ = [8.0] and **w**_**d**,3_ = [1.0] to promote exact matching of the endpoint location, with a smaller emphasis on the accuracy of the tangent matching.

Figure 4 shows one possible motion that accomplishes the defined fruit-picking task. In figs. 4d and e, we demonstrate the underlying activations in the 28 muscular subdomains and the rotation angles of the proximal base that result in the motion visualized in figs. 4a-c. Figure 4a shows the first phase of the motion, while figs. 4b and c visualize the third phase. We omit the visualization of the second phase, i.e., increasing the force applied on the fruit, as it results in very small changes in the geometry of the trunk. The shaded regions in the plots of the muscular activation in fig. 4d and the plots of the angles **ψ**_Z_ and **ψ**_Y_ in fig. 4e delineate the three motion phases. In fig. 5, we show the isolated muscular subdomains during the fruit-picking motion to visualize the deformed muscle geometries with additional clarity. We color the subdomains in fig. 5 according to the magnitudes of their respective muscular activations at multiple time points in both the first and third phase of the motion.

**Figure 4:**
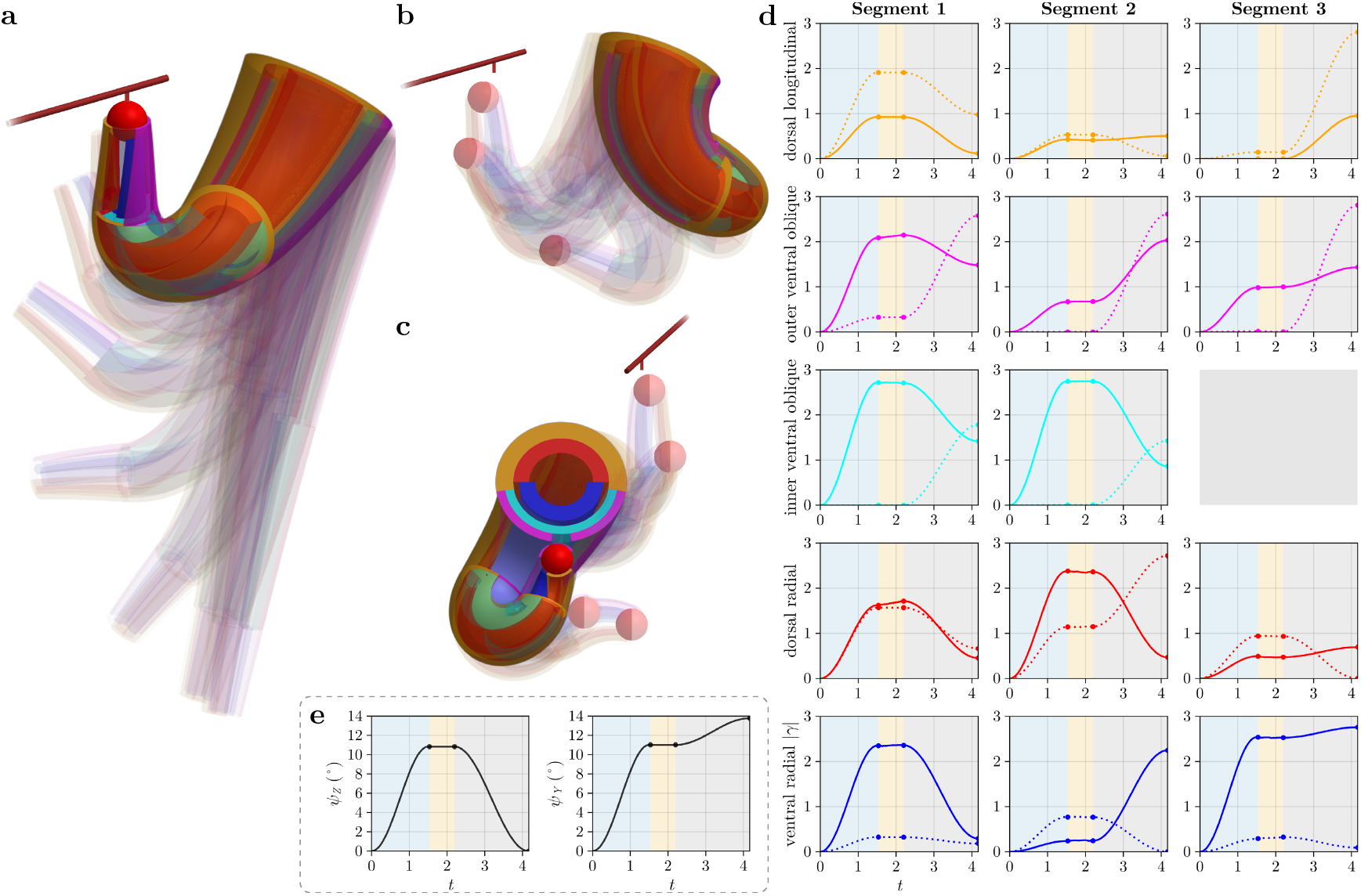
Simulation of an elephant trunk picking and eating a fruit. (a) First phase of the motion in which the trunk reaches the position of the fruit while achieving a prescribed endpoint tangent *–***e**_*Z*_ . (b) Third phase of the motion in which the trunk moves the fruit towards the elephant’s mouth after picking. (c) Back view of the third phase of the motion. (d) Muscular activation magnitudes in 28 muscular subdomains plotted throughout the motion. The rows correspond to different muscle groups and the columns correspond to the three *Z* segments. The solid and dotted lines represent the activations in the right and left trunk, respectively. (e) Rotation angles of the proximal base of the trunk plotted throughout the motion. The light blue, light orange, and gray shaded regions in (d) and (e) correspond to the three motion phases, while the circular markers indicate transition points between consecutive phases.

**Figure 5:**
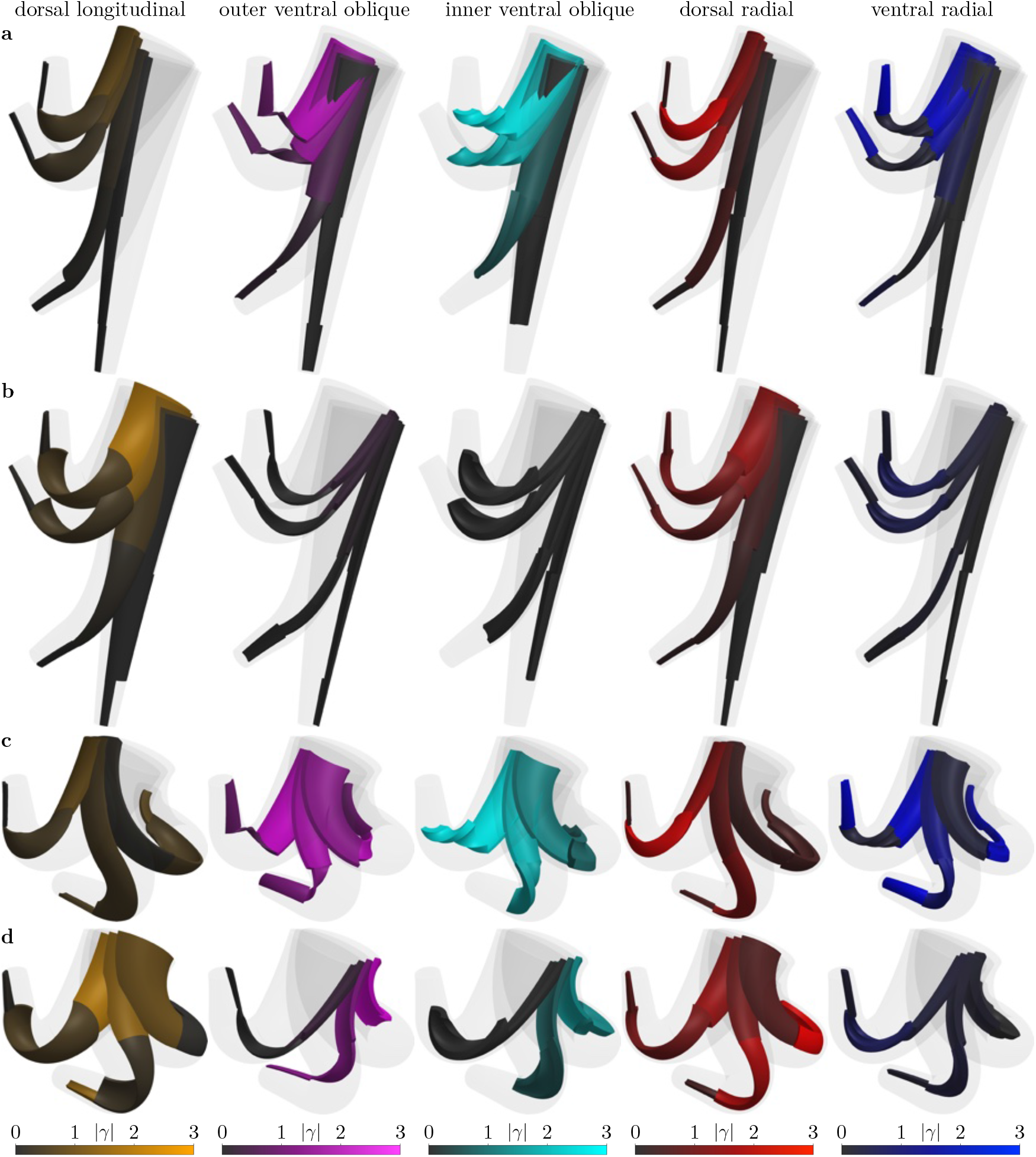
Isolated views of the individual muscle groups during the fruit-picking motion. We color-code the muscular subdomains according to their corresponding activation magnitudes at multiple time points in the (a) right trunk during the first motion phase, (b) left trunk during the first motion phase, (c) right trunk during the third motion phase, and (d) left trunk during the third motion phase. We show four and three consecutive deformations in the first and third phases, respectively. Each column corresponds to one muscle group associated with one color bar legend.

The trunk reaches the fruit location primarily through muscular activation with minor contributions to the motion from the movement of the elephant’s head, as indicated in fig. 4e by the small ranges of rotation angles throughout the motion. In the first phase of the motion, based on figs. 4d and 5a and b, the trunk uses mainly the longitudinal, right oblique, dorsal radial, and right ventral radial muscle groups. The large contraction of the longitudinal muscles in segment 1 bends the entire trunk upwards towards the elevated fruit location, while the similar activations of the right outer and inner oblique muscles roughly cancel the twisting of the trunk in that segment. Despite the large activations of the oblique muscles, the contributions of the longitudinal muscles dominate the bending motion of the first segment. The radial muscles counteract the large axial contraction along the trunk’s centerline due to the longitudinal and oblique muscle activations, which ensures that the trunk’s tip can reach sufficiently far along the *X* direction. The contributions of the radial muscles are small to positive bending around **d**_1_ and negligible to bending around **d**_2_ in the first segment. The deformation of the trunk in segment 2 is densely torsioned thanks to the large activations of inner ventral oblique muscles and small longitudinal muscle contractions. The large twist and torsion in that segment allows the trunk to move closer towards the fruit along the *Y* direction, while orienting itself such that the *Z* = *Z*_2,2_ cross section points upwards in the **d**_3_ = −**e**_Z_ direction towards the fruit. As a result, the short third segment of the trunk does not require much further deformation to reach the fruit location. In addition to the high torsion generated by the oblique muscles in the second segment, the dorsal radial muscles contribute a measurable amount of negative bending around **d**_2_, which facilitates the curling motion in segment 2. Similar to segment 1, the activation of the dorsal radial muscles further counteracts the axial contraction due to the inner ventral oblique muscles. In the third segment, the bending curvatures generated by the outer oblique and radial muscles approximately cancel each other throughout the motion, leading to a roughly straight shape of that segment which successfully reaches the fruit location. In particular, while the twist generated by the right oblique muscle group in segment 3 does not play a functional role in the motion, the activation of this muscle is critical in balancing the bending and axial extension contributions due to the ventral radial muscles.

In the second motion phase, the trunk imparts an increasingly large tensile force on the fruit until reaching a threshold force value that detaches the fruit from the branch. The changes in the underlying activations and base rotations are almost negligible throughout this phase, with very small increases in the muscular activations in the right dorsal radial and right outer ventral oblique groups of the first segment. These small changes are understandable since the activations required to lift the heavy trunk itself in such a highly curled deformed configuration significantly outweigh the activations needed to account for the threshold fruit detachment force of 100 N.

The trunk then proceeds to the third motion phase in which it moves the fruit to the location of the elephant’s mouth. We note that, immediately after picking the fruit, the trunk’s endpoint shifts to a slightly lower position relative to the original location of the fruit. The are two reasons for this shift. First, at the end of the second phase, some of the activations are slightly different than at the beginning of the second phase due to the endpoint tension force and the need to maintain the same endpoint location during activation optimization. Consequently, when the tension force vanishes upon picking the fruit, the new activations result in a different deformed shape of the trunk. Second, in addition to the vanishing of the tension force, picking the fruit results in a change of the boundary conditions whereby the weight of the fruit itself now acts on the endpoint of the trunk. As such, factors both on the activation level and the boundary condition level result in the change in the trunk’s shape immediately after picking the fruit.

In the third phase, the extent of the contributions of each muscle type is highly dependent on the segment in which it resides. To reach the mouth, the trunk seeks to curl the proximal segment towards the body of the elephant. It achieves this curling motion through large contractions, roughly balanced between the right and left trunk, of both the outer and inner ventral oblique muscles in the first segment. The activation of the left longitudinal muscle group also bends the trunk away from the median plane, which provides space for the rest of the trunk, i.e., segments 2 and 3, to reach the mouth without interpenetration. The radial muscles contribute small activations to the deformation of the proximal segment. On the other hand, in the second segment, the radial muscles contract with much higher activations, leading to large contributions to the axial extension of the segment in addition to further bending towards the mouth induced by the oblique muscles. The large activations of the oblique muscles, however, induce overall shortening of the centerline in this segment, despite the large radial contractions. The left oblique muscles assume higher activations compared to the right side of the trunk, which, as indicated in Table 2, provides positive contributions to bending around **d**_1_ from both the outer and inner oblique muscle groups. Given that, by the end of the motion, the second segment curls in a plane close to the horizontal plane, increasing û_1_ prevents the second segment from dropping too far down due to its own weight and that of the remaining third segment. The longitudinal muscles contract to a negligible extent in the second segment, as larger activations would unnecessarily oppose the bending direction contributed by the oblique muscles. By the end of the second segment at *Z* = *Z*_2,2_ and at the end of the motion, the trunk curls by a total angle greater than 180° relative to **d**_3_ at *Z* = 0. As a result, in contrast with the first segment, the dorsal side of the trunk’s third segment moves closer to the mouth than the ventral side. Consequently, large activations of the longitudinal muscles bend the third segment towards the mouth; the ventral radial muscles further amplify the bending in this direction. The additional contractions of the outer ventral oblique muscles introduce some bending in the opposite direction, i.e., they impart negative û_2_ contributions, but these are not sufficient to balance the positive bending due to the longitudinal and ventral radial muscles. The contractions of the muscles in the last segment are asymmetric between the right and left trunk, because the non-zero *Y* component of the tangent **d**_3_ has to transition to zero at *Z* = *L*, so that the distal tangent **d**_3_(*L*) = **d**_3,mouth_ resides in the *XZ*-plane.

### Lifting a log

We proceed with the analysis of the motion consisting of the trunk lifting a cylindrical wooden log. As in the case of the fruit-picking task, we split the motion into three phases: (1) moving the trunk until it just touches the log by wrapping around it from the bottom, (2) establishing full contact until the net force exerted by the trunk on the log equalizes the weight of the log, (3) lifting the log to a prescribed elevated location. We assume that the log is stationary before the lifting phase with its weight supported by an extraneous structure.

We define the first phase as

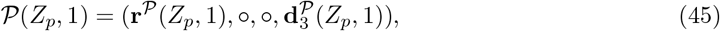

for *Z*_p_ = *Z*_**w**_ + (*p* − 1)(*L* − *Z*_**w**_)/(*N*_p_ − 1), *p* ∈ {1,…, *N*_p_}, where

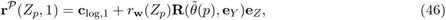

are the points in the discrete log-wrapping trajectory for the *Z* ∈ [*Z*_**w**_, *L*] portion of the trunk’s centerline before lifting the log, **c**_log,1_ is the initial center point of the log located in its *XZ* symmetry plane, 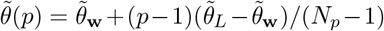 are the discrete angles defining the wrapping trajectory with 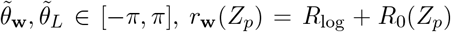 is the distance from the center of the log to the wrapping trajectory at *Z*_p_ that accounts for the tapering profile *R*_0_(*Z*) in defining the contact curve, and *R*_log_ is the radius of the cylindrical log. We assume that the wrapping trajectory **r**^*P*^ (*Z*_p_, 1) results from establishing a one-dimensional contact curve with the log at the intersection of the median plane with the outer surface of the ventral trunk. We define the desired tangents in the wrapping trajectory as

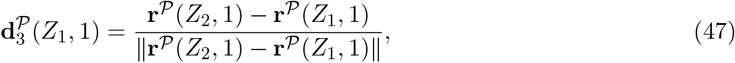

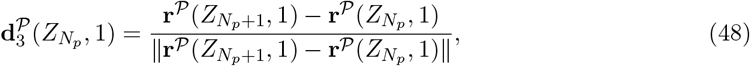

where 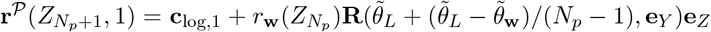 is a fictitious trajectory point extrapolated beyond 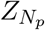. For *p* ∈ {2,…, *N*_p_ − 1}, we set 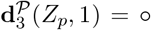, since the entry and exit tangents 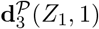 and 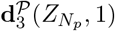 alone provide sufficient definition of the wrapping trajectory. In the first phase, the trunk experiences gravitational forces only, i.e., **N** = **W** = **L** = **0**.

Since, in the second phase, the trunk develops contact with the log until reaching a sufficient lifting force, the function 𝒫 remains the same as in the first phase. In particular, 𝒫(*Z*_*p*_, 2) = 𝒫 (*Z*_*p*_, 1) because, for simplicity, we consider only one time point in the second phase and construct the intermediate motion through interpolation. We found that interpolation results in satisfactory wrapping geometry around the log without solving explicit optimization problems at intermediate loading conditions. In this phase, the weight of the log causes a loading **W** = *W*_log_**e**_Z_ on the trunk, while **N** = **L** = **0** due to the absence of any point loads and since the contact region lies in the *XZ* symmetry plane. Assuming a uniform volumetric density *ρ*_log_ of the cylindrical log, we have 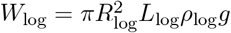.

In the third phase, the trunk lifts the log along a given path,

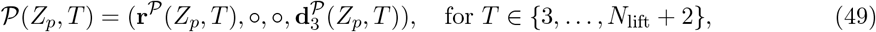

where *Z*_p_ and *p* are the same as in the first two phases, and

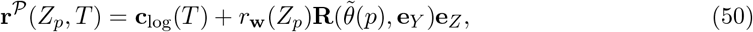

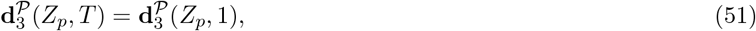

where

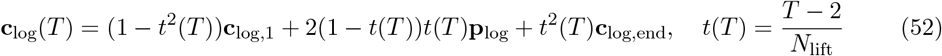

outlines the discrete motion of the log’s center along a quadratic Bézier curve path defined by the points **c**_log,1_, **p**_log_, and **c**_log,end_. We emphasize that the path **c**_log_(*T*) of the log’s center omits the start point **c**_log,1_, i.e., *t*(3) = 1*/N*_lift_ > 0, since **c**_log,1_ generates the same wrapping trajectory as **r**^*𝒫*^ (*Z*_p_, 1) and **r**^*𝒫*^ (*Z*_p_, 2) in the first two phases. In the third phase, the loading conditions are the same as in the second phase, i.e., **W** = *W*_log_**e**_Z_, and **N** = **L** = **0**.

Here we set the parameters of the motion and loading as: *Z*_**w**_ = *Z*_1,3_ − *L*/20, *N*_p_ = 6, **c**_log,1_ = **r**(*L*; **0**) + [−0.4 m, 0.0 m, −0.45 m], *R*_log_ = *L*/24, 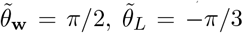, **p**_log_ = **c**_log,1_ + [0.0 m, 0.0 m, −0.4 m], **c**_log,end_ = **c**_log,1_ + [0.2 m, 0.0 m, −0.6 m], *L*_log_ = *L*/2, *ρ*_log_ = 1009 kg/m^3^, *t*_0_ = 0, *t*_1_ = 1.0, *t*_2_ = 1.4, *t*_*i*_ = 1.4+ (*i* − 2)*/N*_lift_ for *i* ∈ {3,…, *N*_lift_ + 2}, *N*_lift_ = 8. The durations of the three motion phases are then 1.0, 0.4, and 1.0, respectively. We use cosine interpolation for ℐ_0_ and ℐ_1_, and cubic Hermite spline interpolation for ℐ_*i*_, *i* ∈ {2,…, *N*_lift_ + 1}. In the optimization problem, |*γ*|_max_ = 3.5, **w**_**r**_ = [7.0, 1.0, 1.0, 1.0, 1.0, 5.0], and **w**_**d**,3_ = [1.0, ○, ○, ○, ○, 1.0]. We set the weights associated with the centerline locations at *Z*_1_ and 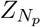 to larger values to ensure satisfactory matching of the entry and exit points in the wrapping trajectory. We found this distribution of weights to be the most effective at facilitating convergence of the optimization scheme to a desirable optimum for the evaluated motion and loading conditions.

In fig. 6, we show the trunk motion that follows the three phases of the log lifting task defined by 𝒫. Figures 6c and d show the magnitudes of activations in the 28 muscular subdomains and the rotations **ψ**_Z_ and **ψ**_Y_ of the trunk base as functions of time *t* throughout the motion visualized in figs. 6a and b. As in the case of the fruit-picking motion, we omit the visualization of the second phase, since, during contact development between *T* = 1 and *T* = 2, the changes in the centerline of the deformed configuration are negligible. The three motion phases correspond to the three consecutive shaded regions in the plots of the muscular activations and base rotations in figs. 6c and d. To more clearly depict the evolution of the muscular activations during the lifting of the log, we show in fig. 7, as before, the isolated muscular subdomains color-coded according to their activations during the third motion phase.

**Figure 6:**
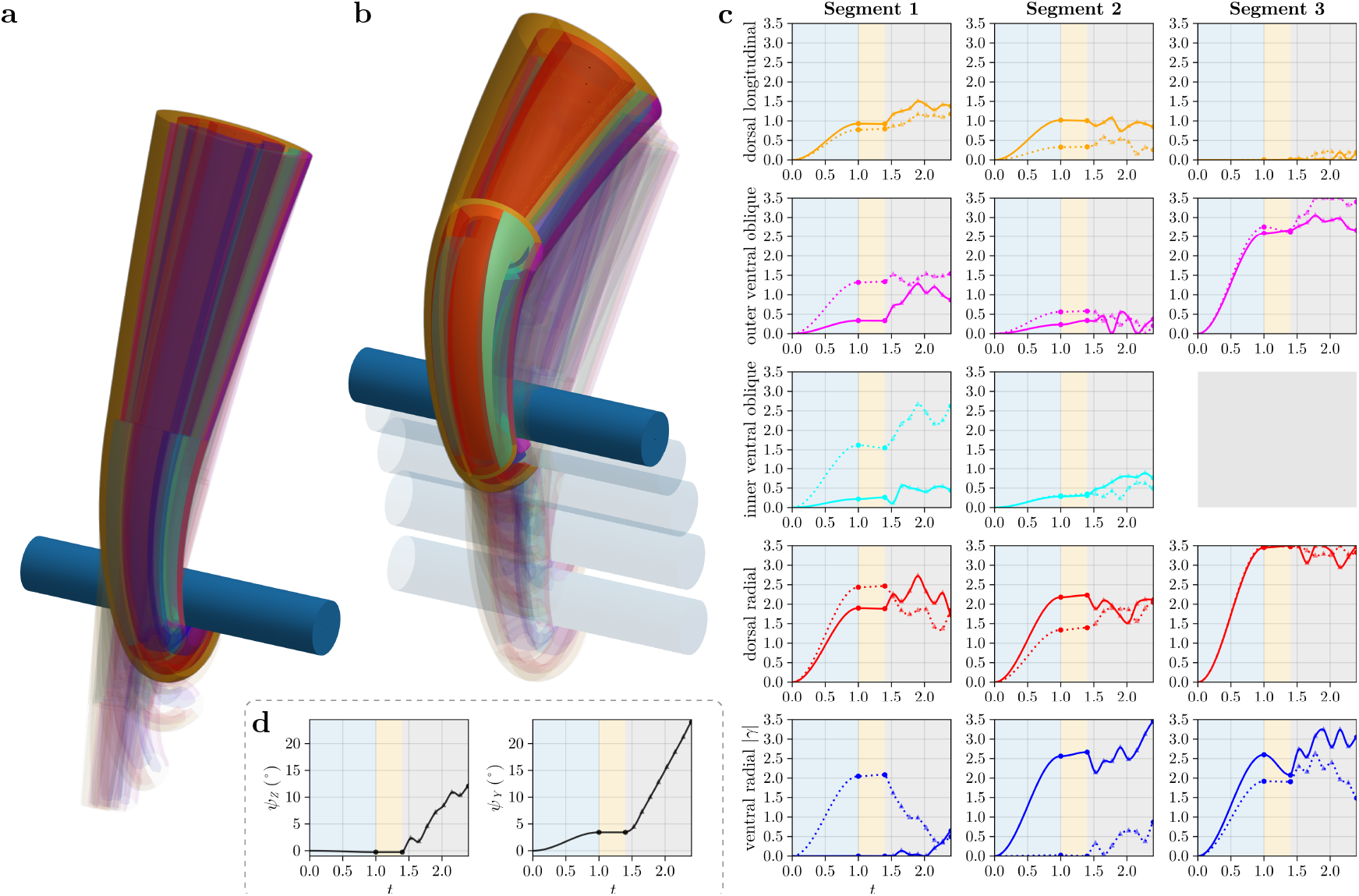
Simulation of an elephant trunk lifting a wooden log. (a) First motion phase during which the trunk curls to achieve a prescribed wrapping shape **r**^*𝒫*^ (*Z*_*p*_, 1) in the interval *Z* ∈ [0.8*L, L*]. The trunk then establishes contact with the log in the second phase while seeking to preserve the same wrapping trajectorχ. (b) Third motion phase in which the trunk lifts the log along a prescribed path defined bχ the movement of the log’s center **c**_log_. (c) Muscular activation magnitudes for the 28 muscular subdomains in the right and left trunk plotted throughout the motion as solid and dotted lines, respectively. (d) Rotation angles of the proximal base of the trunk plotted throughout the motion. In (c) and (d), the light blue, light orange, and gray shaded regions correspond to the three motion phases, the circular markers indicate transition points between phases, and the semi-transparent triangular markers indicate the eight time points *T* ∈ {3,…, 10} along the lifting path.

**Figure 7:**
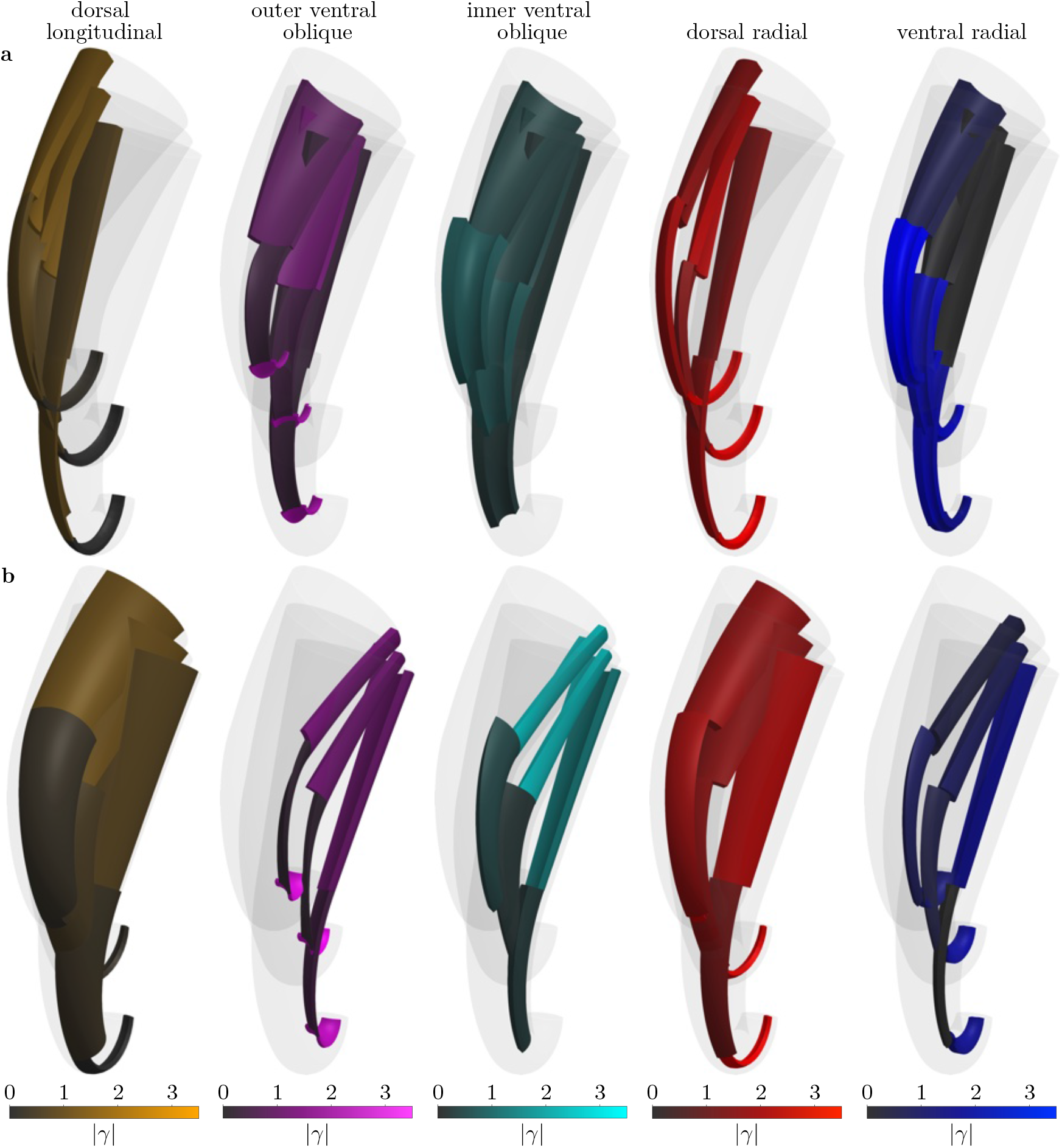
Isolated views of the individual muscle groups and their activations during the log lifting task. We color-code the muscular subdomains according to their corresponding activation magnitudes at three time points in the (a) right and (b) left trunk during the third motion phase. Each column corresponds to one muscle group associated with one color bar legend.

We begin our interpretation of the results with several general observations that hold throughout the motion. In contrast to the fruit-picking motion, the rotation of the trunk’s proximal base is more significant in the log lifting scenario. For more restrictive feasible sets imposed on **ψ**_Z_ and **ψ**_Y_ , the optimization method could not find optima with sufficiently small objective function values, which suggests that non-negligible rotation of the elephant’s head is inherent in this motion, given the chosen constraints on the activations **Γ**; see Appendix B. Throughout the motion, the trunk activates the longitudinal muscles to the smallest extent out of all the muscle groups. Specifically, the solution of the inverse problem yields small longitudinal muscle activations since they can only provide positive contributions to û_2_, while lifting the log requires a curling motion towards the log which translates to negative û_2_. Measurable activations are still present, however, in the first segment of the dorsal longitudinal muscles, which contributes an overall lifting action to the rest of the trunk.

We observe moderate-to-high activations in the oblique muscle groups, with the largest contractions in the outer ventral oblique muscles of the third segment. The third segment *Z* ∈ [*Z*_1,3_, *Z*_2,3_] is primarily responsible for wrapping around the log since the wrapping trajectory uses *Z* ∈ [*Z*_1,3_ − *L*/20, *Z*_2,3_]. Segment 3 achieves the wrapping shape through large contractions of the ventral oblique muscles that provide negative bending contributions to û_2_, which translates to bending around the log. Further, the trunk ensures minimal torsion in the third segment—which is a property of the desired wrapping trajectory—through approximately symmetric contractions between the respective right and left muscles in all muscle groups. Interestingly, large activations in the left inner ventral oblique muscles of the first segment induce significant positive twist, which is visually evident in the rotated proximal cross section of the second segment. The twisting of the first segment likely ensures a more desirable rotation of the second segment, so that the trunk establishes the trajectory entry condition 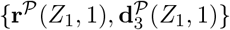 by the end of the first phase and maintains it throughout the third phase with the additional contact loading.

Among all muscle groups, the radial muscles exhibit the highest activations. In all segments, the large radial activations provide extensile contributions to the trunk that counteract the centerline contraction due to other muscles which, in turn, enables reaching the distant log while still matching the wrapping trajectory. In particular, high radial contractions are the most predominant in the third segment, where the large activations of the outer ventral oblique muscles alone would otherwise make it impossible to match the wrapping trajectory due to centerline contraction. Concretely, large shortening of the third segment would cause an incomplete wrapping geometry, likely leading to the log falling. Noteworthy is also the right-left asymmetry in the activations of the ventral radial muscles of the first two segments. The contractions of the right ventral radial muscles in the second segment provide measurable positive contributions to the bending curvature û_1_ with little to no opposing contributions from the corresponding left group. Further, the left ventral radial muscles in the first segment contribute negative bending to û_1_ in the first and second phase, with no opposing activation in the right group. In the third phase, the asymmetry in the activations of the ventral radial muscles of the first segment reduces as a result of a significant decrease in the activations of the left ventral radial group. The decrease in the activation in that radial subdomain is likely since the lifting motion brings the log to a higher position, requiring smaller extensile contributions from the group.

Perhaps the most striking feature of the extracted muscular activation profiles is the high degree of non-monotonicity of the activation magnitudes in the third phase. While the optimization at multiple intermediate points during the lifting part of the motion is certainly a factor in producing this non-monotonic behavior, there are other critical mechanisms that yield such variation of muscular activations in time.

First are the constraints imposed on the muscular activations, the trunk base rotation angles, and the activated centerline extension. The penalty function 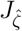 effectively limits the trunk extension to be in the range 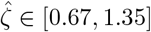. The activation magnitudes cannot exceed |*γ*|_max_ = 3.5. Further, both the trunk base angles **ψ**_Z_, **ψ**_Y_ and the activation magnitudes obey additional restrictions of the form *x*_*i*_(*T*) ∈ [*x*_*i*,min_(*T*), *x*_*i*,max_(*T*)]; see Appendix B. These constraints limit the space of accessible configurations. As a result, if one of the quantities seeks to cross into the infeasible region to achieve the desired configuration properties, then all muscles need to search for another optimal arrangement, so that all activations ultimately stay within the constraint boundaries. In the limit of a continuous-time problem with the number of optimization runs *T*_max_ approaching infinity, the global optimum can shift abruptly between consecutive time points due to the constraints despite a designation P that is continuous in time. In other words, even though the sequence of consecutive optima (**x**^*\**^(*T*), **x**^*\**^(*T* + 1),…) often exhibits small changes between successive **x**^*\**^, encountering the boundary can force the optimization scheme to search in an entirely different area of the objective function landscape to identify the global optimum. How far the potentially more desirable area of the objective function can be relative to the original area depends on the choice of the additional constraint *x*_*i*_(*T*) ∈ [*x*_*i*,min_(*T*), *x*_*i*,max_(*T*)]. As an example, the activation in the dorsal radial muscles in segment 3 reaches the |γ|_max_ = 3.5 boundary already by the start of the third phase. To move the log in an upward motion while still satisfying the constraints, the activations in the dorsal radial muscles of the third segment decrease to move away from the |γ|_max_ boundary, while the activations of other muscle groups adjust in either positive or negative directions. The new global minimum lies in an area more removed from the feasible set boundary. A similar process repeats once the dorsal radial muscles in segment 3 reach the |γ|_max_ boundary again in the middle of the third phase.

Second, the variation in muscular activations during the lifting phase is also due to a phenomenon unrelated to constraining the design variables. In particular, the objective function landscape changes as a function of time since every optimization problem defines a different function *J*_T_ constructed based on 𝒫 at a given *T*. If we again assume a continuous time domain for 𝒫 with an infinite number of consecutive optimization problems, the local minimum locations vary continuously in time, while the global minimum locations can change discontinuously due to the evolving objective function landscape. As such, in the discrete time setting, abrupt changes between the successive optima **x**^*\**^(*T*) and **x**^*\**^(*T* + 1) can occur purely because points **x** at *T* + 1 that have lower objective function values than the previous optimum **x**^*\**^(*T*) emerge in other regions of the feasible set. The previously mentioned additional constraints *x*_*i*_(*T*) ∈ [*x*_*i*,min_(*T*), *x*_*i*,max_(*T*)] on the muscular activations and the trunk base angles prevent such jumps across the design space from becoming excessively distant.

The general process of reorganization of the fibrillar activation throughout the trunk could likely occur in the real elephant trunk as well. However, it would presumably be a smoother process, as the activation rearrangements would most likely occur in anticipation of the boundary being approached, rather than after reaching the boundary. That is, an elephant might have some intuitive foresight into which solution branch of the inverse problem to pursue before lifting the log, so that it does not need to frequently reorganize the activation distribution throughout the trunk. In contrast, the optimization process in our model responds to constraint boundaries only upon reaching them, as future points along 𝒫 do not inform the optimization at the current time point.

### Asymmetric lifting of a log

Finally, we investigate the effect of asymmetric lifting on the underlying muscular activations. The setup of this motion is almost identical to the symmetric log-lifting scenario with the main exception being that the center of the wrapping trajectory is no longer aligned with the center of the cylindrical log. In particular, the center of the wrapping trajectory is at a distance *d*_offset_ from the center of the log in the **e**_Y_ direction. While the definition of 𝒫 remains the same as in the symmetric case, the offset wrapping trajectory gives rise to a couple

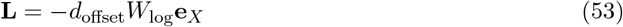

imposed on the trunk along *Z* ∈ [*Z*_**w**_, *L*] in the second and third phases of the motion. The equal and opposite couple −**L** ensures moment balance for the log during these phases. To emphasize the effect of the resulting couple, we set *d*_offset_ = 0.3*L* and elongate the cylindrical log length to *L*_log_ = 0.75*L*. For the optimization procedure, the maximum permissible activation magnitude is |γ|_max_ = 3.5 as in the symmetric lifting scenario. We decrease the weights for the tangent director to **w**_**d**,3_ = [0.5, ○, ○, ○, ○, 0.25] because, as long as the distal end of the trunk matches the wrapping trajectory **r**^*𝒫*^ , we do not seek to penalize tilting resulting from the external couple.

Figures 8 and 9 show the resulting motion of the elephant trunk lifting the log with off-center contact. We generate the plots and visualizations in these figures equivalently to the respective fig. 6 and 7 for the symmetric lifting case. The motion, the activations, and the trunk base angles in the first phase shown in figs. 8 are identical to those in fig. 6 since the loading conditions and the motion specification 𝒫 is the same in both cases at *T* = 1. The asymmetric lifting scenario deviates from the symmetric case in the second phase, during which the trunk develops contact with the log and the external couple **L** emerges. We observe that the couple and the muscular activations that achieve the wrapping trajectory result in tilting of the trunk towards +**e**_Y_ which acts as leverage against the applied couple. To achieve a satisfactory matching of the lifting motion, we permit larger deviation ranges for the trunk base angles at each *T* relative to the solution at *T* − 1; see for the discussion of feasible sets as functions of *T*. Consequently, the contributions of the rotations of the trunk’s proximal base are the most significant in this motion with **ψ**_Y_ reaching about 30.5°. Nevertheless, the deformation due to muscular activity still largely dominates the qualities of the motion that lead to the lifting of the log. Similar to the symmetric case, the muscular activations in the third phase are non-monotonic in time with moderately large changes between subsequent time points. The explanation of non-monotonicity for the symmetric lifting motion applies to the asymmetric scenario as well. The activation rearrangement behavior observed in the symmetric case is also present in this motion, as evidenced by the activation curves in the dorsal radial muscles of the third segment. However, for some muscle groups in the third phase, the changes in activation magnitudes between consecutive time points are larger compared to the symmetric case since the feasible activation ranges used in this motion are wider. As in the case of the trunk base angles, we found that less restrictive constraints were necessary to yield sufficiently low objective values for a reasonable number of optimization runs.

**Figure 8:**
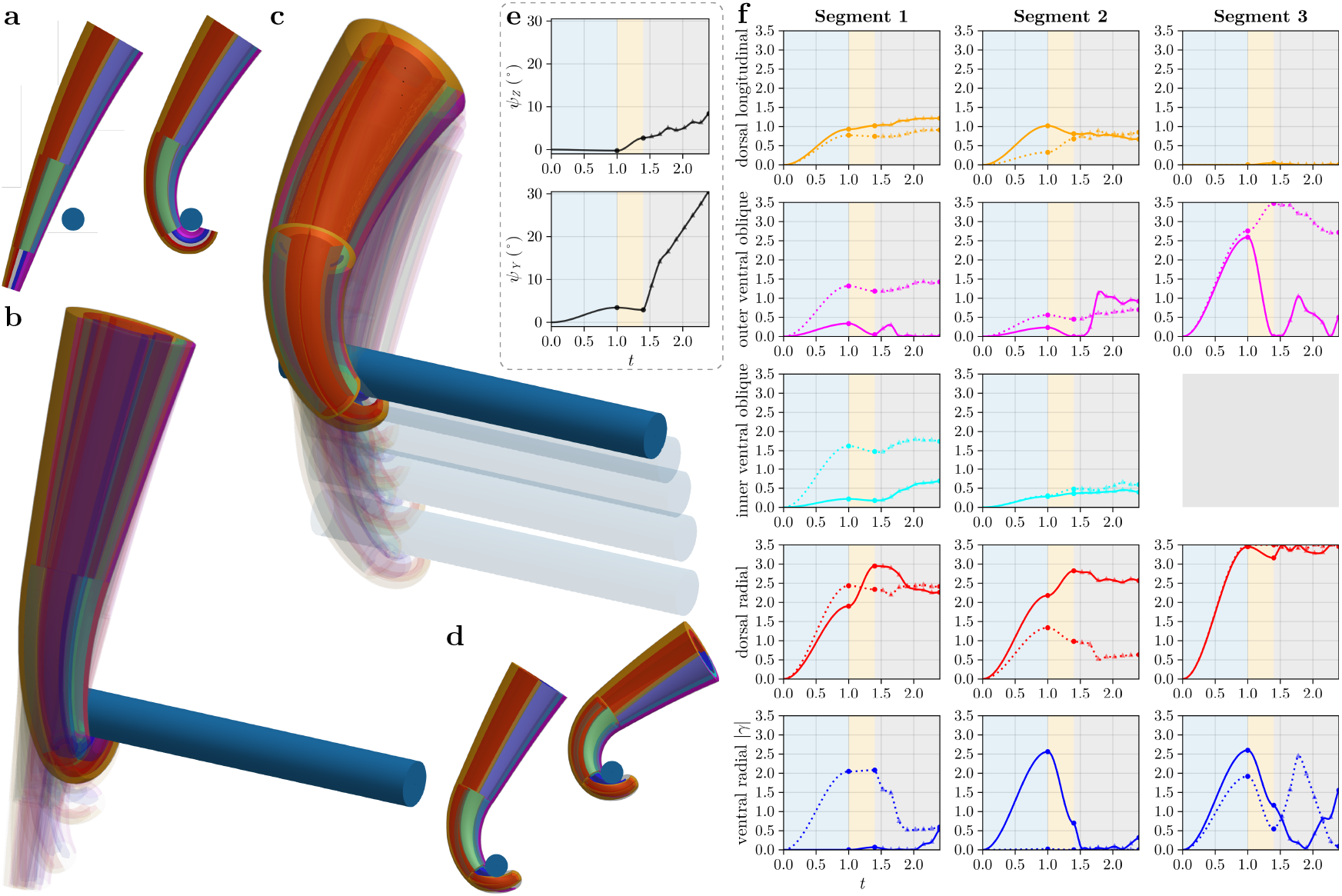
Asymmetric lifting of a wooden log. (a) Side view of the first and last configurations in the first motion phase. (b) First motion phase in which the trunk curls around the offset wrapping trajectory without any contact forces or couples. (c) Third motion phase in which the trunk lifts the log while wrapping around it according to an offset trajectory, which gives rise to a non-zero couple in addition to the distributed load due to the weight of the log. (d) Side view of the first and last configurations in the third motion phase. (e) Rotation angles of the trunk’s proximal base plotted throughout the motion. (f) Muscular activation magnitudes for the 28 muscular subdomains plotted throughout the motion. We visualize the plots in (e) and (f) in the same manner as in figs. 6d and c.

**Figure 9:**
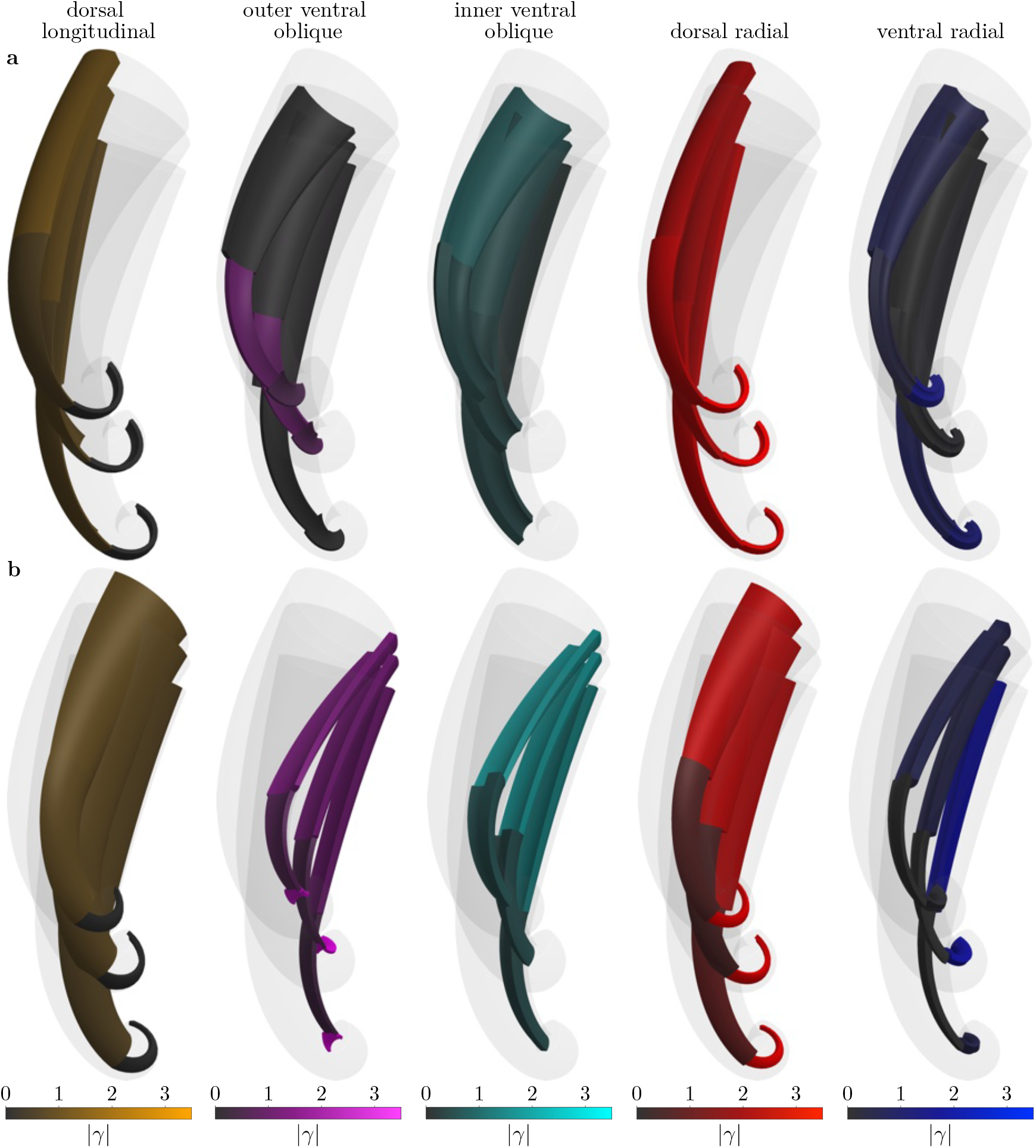
Isolated views of the individual muscle groups and their activations during asymmetric log lifting. As in fig. 7, the color-coding corresponds to the activation magnitudes in (a) right and (b) left trunk during the third motion phase.

One of the primary differences between the symmetric and asymmetric lifting cases is the effect of the applied external couple on the underlying muscular activations. In addition to the relatively large range swept by the angle **ψ**_Y_ , we note the significant change in the muscular activations in the second phase relative to the first phase. The couple considerably shifts the desirable minimum in the design space relative to the couple-free minimum, which requires large changes in the muscular activations to achieve the desired wrapping trajectory with sufficient accuracy. As such, for the optimization problem at *T* = 2, we permit values of muscular activations in the entire range *x*_*i*_[−|γ|_max_, 0], *i* ∈ {1,…, 28}, irrespective of the optimization result at *T* = 1; see Appendix B. By doing so, we allow a minimum **x**^*\**^(*T* = 2) to emerge with a sufficiently small *J*_2_.

Importantly, the addition of the external couple results in extensive asymmetries in the activations between the right and left trunk in the third phase. Since **L** acts along the −**e**_X_ axis, the trunk needs to oppose the couple with differential activation along **e**_Y_ , which translates to vastly different activations required in the right and left muscle groups. In particular, compared to the large activations in the respective opposing muscle group, we observe significantly lower activations in the right outer ventral oblique muscles in segments 1 and 3, right inner ventral oblique muscles in segment 1, and right ventral radial muscles in segment 1. Notable right-left asymmetries are also present in the dorsal radial muscles in segment 2 and ventral radial muscles in segment 3. From a functional standpoint, the mismatch between the right and left activations has the most prominent effect on the generation of twist and torsion by the oblique muscles. For equal activations in the right and left oblique muscle groups, the twist and torsional contributions cancel each other resulting in pure bending. In contrast, the significantly larger activations in the left outer ventral oblique muscles in segment 3 introduce large negative contributions to twist and torsion. Nevertheless, little twist emerges in the first segment, despite much larger activations in the left oblique muscles compared to the right oblique groups. The signs of the twist contributions of the outer and inner ventral oblique muscles are opposite, see Table 2, which leads to cancellation of mechanical effects across different muscle architectures rather than across the two sides of the trunk.

Beyond the asymmetries between the right and left trunk, we note the differences in the overall activation magnitudes between the symmetric and asymmetric lifting motions. Specifically, in the lifting phase, the following activation curves exhibit much lower values as compared to the symmetric scenario: right outer ventral oblique in segments 1 and 3, left inner ventral oblique in segment 1, left dorsal radial in segment 2, and right ventral radial in segments 2 and 3. These lower activation magnitudes might be in part due to higher contributions of the _Y_ angle to the lifting action as compared to the symmetric case.

## 7. Conclusions

We have created a model of the elephant trunk to simulate the biomechanics of the trunk’s soft musculature by using formal continuum-mechanics arguments. The geometrically exact formulation of the elephant trunk as an extensible Kirchhoff rod ensures reliable predictions for large deformations. We used magnetic resonance images [38] to inform the cross-sectional anatomy of the trunk’s musculature throughout its length. By representing the trunk as an active slender structure, the dimensional reduction in the deformation map results in closed-form formulas for the activated curvatures and extension. These analytical expressions provide real-time simulation functionality, whereby the computation of a deformed trunk configuration takes only a fraction of a second on a standard desktop computer. The low computational cost of the model enables efficient solution of the inverse motion problem of predicting the muscular activations required to match a desired deformation.

To solve the inverse motion problems, we used global optimization with an objective function that quantifies the deviations from the desired geometry. The high performance of the model is critical to the employed optimization methodology, as it evaluates a large number of muscular activation candidates to converge to a solution that matches the target deformation. By solving a time series of optimization problems with varying deformation targets, we can construct a quasistatic motion of the trunk that achieves a given physiologically relevant task. Since the motion construction involves solving the inverse problem, we automatically obtain the underlying time series of muscular activations.

Using this method, we evaluated three trunk motions and analyzed the muscular biomechanics governing each motion. In our analysis, we uncovered general principles that might govern the biomechanics of the real elephant trunk, such as: mid-motion rearrangement of muscular activations, amplifying or counterbalancing the mechanical contributions of other muscles, moving the trunk’s proximal base to enlarge the space of achievable function, or exploiting architectural and geometrical asymmetries to either amplify other biomechanical effects or induce asymmetry in the deformation.

Our study has several limitations. First, the automatic optimization method generally requires problem-dependent adjustments to achieve minima with sufficiently low objective function values. Instead, one might reasonably resort to manually searching the activation space for the muscular activations that result in a sufficiently close matching of the prescribed task. However, adjusting the muscular contractions manually is extremely difficult since activating any proximal muscle fibers has a snowball effect on the remaining distal portion of the trunk. Further, the interactions of the longitudinal, oblique, and radial muscles produce complex coupling mechanisms between extension, bending curvatures, and torsion. These couplings are non-trivial to employ or bypass intuitively during manual activation design. Our trunk model and the associated inverse problem solution method provide a feasible pathway to extract the muscular activations for different motions, which enables quantitative discovery of the underlying muscular principles.

Second, our trunk representation uses three segments, each of which employs a single muscular geometry setup informed by the corresponding magnetic resonance image [38]. This piecewise structure of the trunk model is approximate and provides only an averaged biomechanical behavior for each segment. In principle, we could split the trunk into more segments and inform each of them by the corresponding magnetic resonance images. However, since each segment involves at least eight independent muscular activations, doing so would proportionally increase the dimensionality of the design space in the optimization problems which, in turn, could make the computation of the motions infeasible. Alternatively, we could permit non-uniform fibrillar activation in each of the three segments by setting 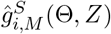 to functions interpolating unknown two-dimensional activation data in each muscle. However, increasing the number of interpolation points would again translate to increased computational cost of solving the inverse problem.

Third, the evaluated motions are quasi-static since the deformed configurations derive from solving the equilibrium boundary value problem at each time point. By considering a non-zero right-hand side in eq. (11), we could also consider the dynamics of the motion, albeit at an increased computational expense and conceptual complexity in the inverse problem solution approach. Fourth, we note that the extracted muscular contractions might exceed the physiological regime, because the constraint |γ|_max_ derives from the empirically smallest feasible set that still enables the construction of a given motion. Future work could entail experimental validation of the model, in which recorded muscular activations and observed deformations of the trunk would quantitatively contextualize the pre-strain measure *ĝ* in a physiological setting. Nevertheless, we emphasize that the trunk’s physiology still directly informs the permissible activations in our model since we prohibit activations that result in leaving the physiologically-informed range 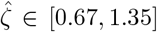 for the trunk’s contraction or extension.

Finally, the muscular activation rearrangement phenomenon discussed in this study could be a manifestation of a limitation in our optimization methodology. In particular, in our optimization approach, we solve each optimization problem at a given time point without considering the potential biomechanical requirements of future time points, given the imposed constraints. To simulate the intent and foresight of the elephant seeking to perform a particular task, each optimization problem should, through some quantitative planning process, incorporate the context of the remaining part of the motion.

Our work provides quantitative insights into the biomechanical intricacies of the elephant trunk through reduced-order modeling. This complex and fascinating structure remains a subject worthy of further investigation. In tackling the formidable challenge of trunk mechanics, we also come closer to completing our essential understanding of other active slender structures in nature. Beyond extending the fundamental science, a thorough understanding of these biological marvels could prove invaluable for inspiring the material science of flexible structures, the design of soft robots, and the creation of flexible prosthesis and assist devices.

## Acknowledgments

Bartosz Kaczmarski acknowledges the Burt and Deedee McMurtry Stanford Graduate Fellowship in Science and Engineering. Zéphyr Goriely gratefully acknowledges the hospitality of the “Save the Elephants” organization at Samburu National Reserve as well as the help of Prof. Fritz Vollrath. Ellen Kuhl acknowledges support through the NSF Grant CMMI 231818 and the ERC Advanced Grant 101141626.

## Appendix A. Activated extension and curvatures

Here we give the explicit form of the activation terms *H*_j_ in eq. (24). Recall that, based on the split of the domain into 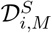 and the activation formulas in eq. (6), the *H*_j_ numerator terms of the activated extension and curvatures become piecewise functions in the three *Z*-segments such that

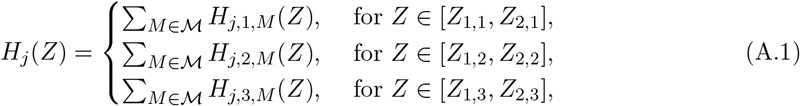

where *j* ∈ {0, 1, 2, 3}, ℳ = {dl, ovo, ivo, dr, vr}. Assuming a homogeneous Young’s modulus *E* and Poisson’s ratio *ν*, we have

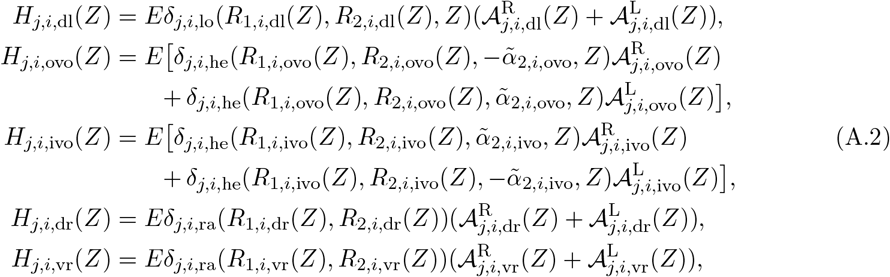

for *j* ∈ {0, 1, 2, 3}, *i* ∈ {1, 2, 3}, where

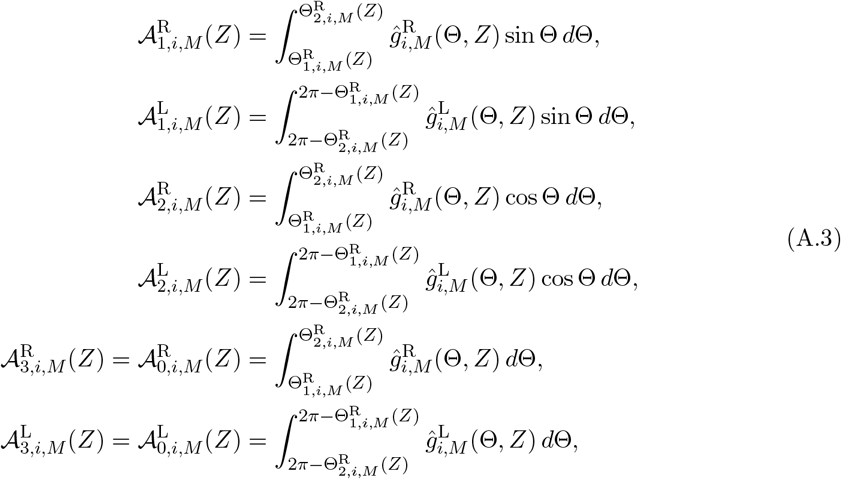

for *i* ∈ {1, 2, 3}, *M* ∈ ℳ, and, assuming that *ϕ* (*R, Z*) ∈ (0, *π*/2],

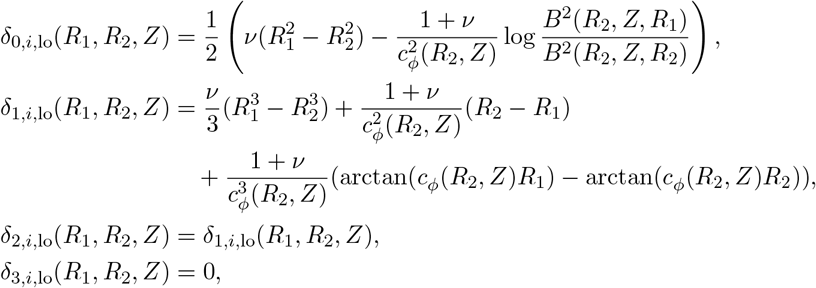

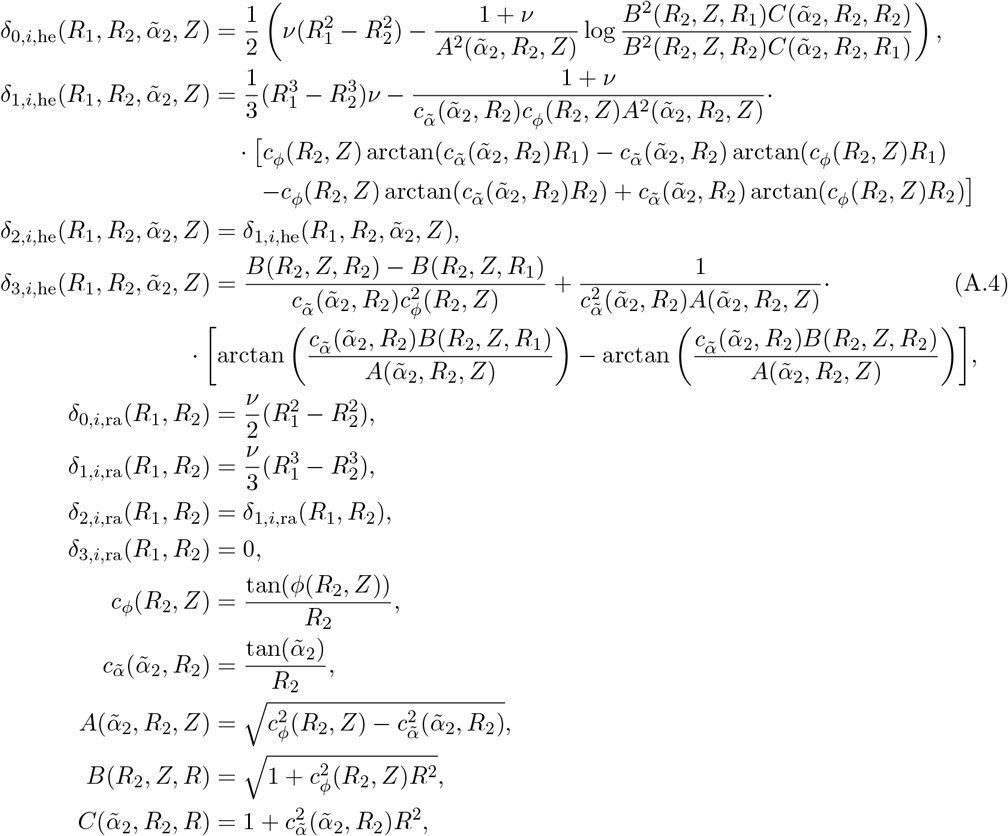

for *i* ∈ {1, 2, 3}, and *Z* ∈ [*Z*_1,*i*_, *Z*_2,*i*_]. The 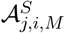 quantities are the integrals of the activation in muscle *M* on side S that enter the corresponding *j*-th curvature component in the *i*-th *Z*-segment. The *δ* functions are prefactors that depend on the geometry and fiber architecture of a given muscular subdomain, and the Poisson’s ratio of the trunk. The designations ‘lo’, ‘he’, and ‘ra’ in the subscripts of the *δ* functions dictate the different prefactor forms for the longitudinal, helical, and radial fiber architectures, respectively. Splitting the *δ* quantities into these three groups simplifies the form of eq. (A.2).

## Appendix B. Feasible sets and initial guesses

Constructing sufficiently restrictive feasible sets is critical to achieve suitable computational performance in solving the optimization problems. The functional landscape of J_T_ at every time point *T* generally exhibits numerous local minima, so identifying the global minimum over ℝ^30^ can be costly for a large feasible set. At the same time, restricting the feasible set can inadvertently remove minima with lower objective function values, which correspond to better matching of the desired configuration properties. As such, choosing an appropriate feasible set size is critical for locating a minimum with a low objective function value while maintaining a reasonable computational effort.

Our approach for constructing the feasible sets is largely heuristic and depends on the behaviour of a chosen optimization scheme for a particular trunk motion. In all cases, we use computational information either from previous time points or from auxiliary optimization results to inform the feasible set or the initial guess in each optimization problem. Given a reference minimum 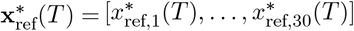 for a time *T* obtained from auxiliary optimization or optimization at a previous time point, we build neighborhood sets for the muscular activation and the trunk base angles around that reference point as the Cartesian products

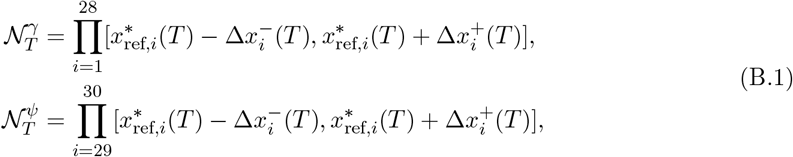

Where 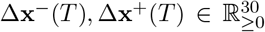 are vectors of admissible deviations from the reference point at a time *T*. Together with the contractile activation constraint *x*_*i*_ ≤ 0, *i* ∈ {1,…, 28}, and the maximum activation magnitude constraint |*x*_*i*_|≤|*γ*|_max_, *i* ∈ {1,…, 28}, the feasible sets become

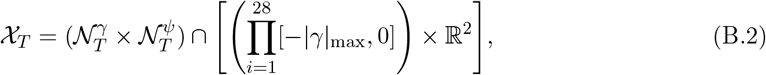

for *T* ∈ {1,…, *T*_max_}. We note that the penalty function 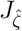 further restricts the viable regions of each feasible set during optimization.

Since each evaluated motion poses different computational challenges and exhibits a distinct behavior in the optimization process, we use different approaches to define 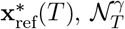,and 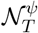 depending on the motion. Importantly, computing the activated configuration **ℬ** is equivalent to solving an initial value problem, which is significantly faster than solving the boundary value problem for the deformed configuration **ℬ**_d_. We can compute the activated extension and curvatures using eq. (5) and then rapidly integrate the kinematics equations to obtain the activated centreline 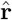 and directors 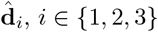 We refer to an optimization problem without external loading and with the substitutions 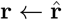 and 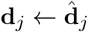 in J_*T*_ in eq. (41) as an *auxiliary* optimization problem. We call the original form of eq. (41) the *main* optimization problem. We denote the minima obtained from the auxiliary and main problems as 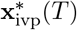 and **x**^*\**^(*T*), respectively. Wherever applicable, 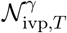 and 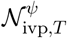 are the sets used in the auxiliary problem, and 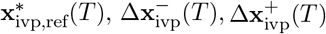 are the auxiliary quantities defined as in eq. (B.1).

Below, we include the specifics of the feasible set construction and the initial guess selection methodology. For some optimization problems, we define the 𝒩-sets explicitly and without a reference point. In all motions, we seek to limit the potential functional dominance of the trunk base rotation over the muscular activations, since the biomechanical effects of muscular contractions constitute the primary subject of this study. As a result, our choices of 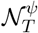 are appropriately more restrictive. We emphasize that the methods and parameters described here are not the only ones that yield satisfactory motion results. Nonetheless, in our heuristic investigation, the choices provided below generated the most desirable motion features out of numerous other choices tested.

### Picking a fruit

In the first phase, we use the optimum 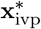 (1) obtained from the auxiliary problem at *T* = 1 only as an initial guess for the main optimization problem and not as a reference point. The feasible set χ _1_ follows simply from the condition *x*_*i*_ ∈ [−|*γ*|_max_,0], for *i* ∈ {1,…, 28}, and an additional restriction *ψ*_Z_, *ψ*_Y_ ∈ [−*π*/16, *π*/16] on the trunk base angles. Then, 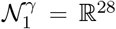 and 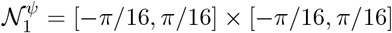 which yields X_1_ through eq. (B.2). We use the same feasible set to obtain the auxiliary 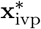 (1) itself. Throughout the motion, we set |*γ*|_max_ = 3.0.

In the second phase, the reference points are not the solutions to the auxiliary problems, but rather the solutions to the main optimization problems at the previous time points. Specifically, we set (*T*) = 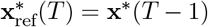,for *T* ∈ {2,…, *N*_pick_ + 1}, where **x**^*\**^(*T* − 1) is the solution to the main optimization problem at *T* — 1. We then define the deviations 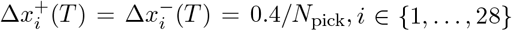, and 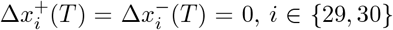 ,The optimum **x**^*\**^(*T* − 1) also serves as the initial guess for the main optimization problem at time *T*.

At *T* = *N*_pick_ +2, we use the optimum 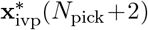 from the auxiliary optimization problem as an initial guess for the main optimization and to build the muscular activation set 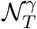. The auxiliary problem that yields 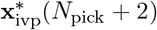 uses a feasible set with 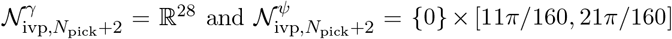. For the main problem, we prescribe 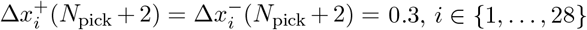, and 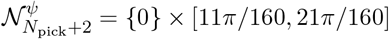.

### Lifting a log

Throughout this motion, we set |*γ*|_max_ = 3.5. In the first phase, we use 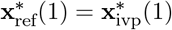 , where the auxiliary problem uses a feasible set defined by 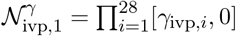 where

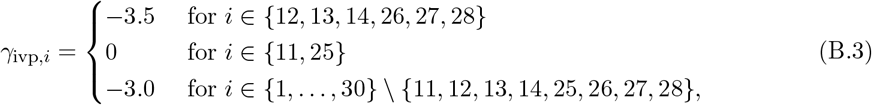

and 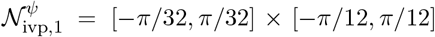 For the main optimization problem, 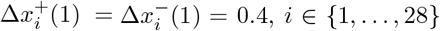 ,*i* ∈ {1,…, 28}, and 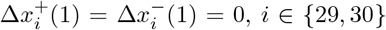,*i* ∈ {29, 30}. The auxiliary 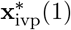 is the initial guess for the main optimization problem.

Similarly, in both the second and third phases, we use the auxiliary problem optima as the reference points, i.e., 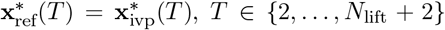. Each 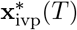 is also the initial guess for the main optimization problem at time *T*. For a given auxiliary problem at time *T*, we use the previous auxiliary optimum as the initial guess and the auxiliary reference point, so that 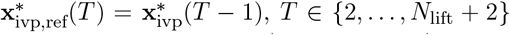, *T* ∈ {2,…, *N*_lift_ + 2}. We set 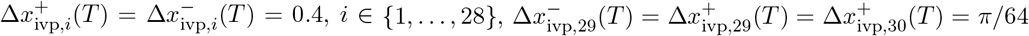 and 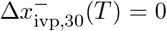 to build 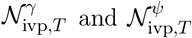 . Then, for the feasible sets in the main optimization problems we choose 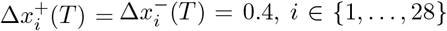 , *i* ∈ {1,…, 28}, and restrict the trunk base angles in the main problem to follow the obtained auxiliary optima exactly, i.e., 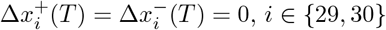, *i* ∈ {29, 30}.

### Asymmetric lifting of a log

Similar to symmetric lifting, the asymmetric lifting motion includes the activation magnitude constraint |*γ*|_max_ = 3.5. In the first phase, the motion is equivalent to the symmetric case since the trunk only experiences gravitational forces due to its own weight. As such, the main problem optimum **x**^*\**^(1) in this motion is the same as **x**^*\**^(1) obtained for symmetric lifting.

From the second phase onward, the trunk experiences the additional external couple. In contrast to the symmetric scenario, the couple generally causes the main problem optimum to lie far from the auxiliary problem optimum. As a result, it is less effective to use the auxiliary optima as reference points for the main optimization problem. Instead, we proceed with sequential solution of the main optimization problems for *T* ∈ {3,…, *N*_lift_ + 2} by using the previous optima **x**^*\**^(*T* − 1) as reference points. That is, we set 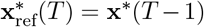,for *T* ∈ {3,…, *N*_lift_ + 2}. To obtain a valid starting reference point 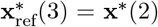 for the problem at *T* = 3, we first use **x**^*\**^(1) as the initial guess for the problem at *T* = 2. The optimization at *T* = 2 utilizes a non-restrictive 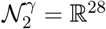 for the muscular activations and 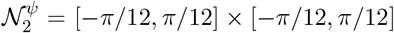 for the trunk base angles. For the feasible sets in subsequent optimization problems, we choose:

- 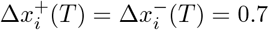,for *i* ∈ {1,…, 28} and *T* ∈ {3, 4, 6, 7, 8, 9, 10},
- 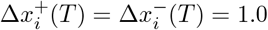, for *i* ∈ {1,…, 28} and *T* = 5,
- 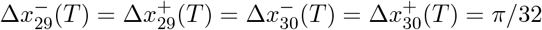, for *T* ∈ {3, 4},
- 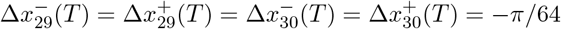,for *T* ∈ {5, 6, 7, 8, 9, 10},

with *N*_lift_ = 8 used in the third phase. The initial guesses for the optimization problems at *T* ∈ {3,…, *N*_lift_ +2} are the optima **x**^*\**^(*T* −1) at the respective previous time points with additional random perturbations 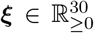 added to each **x**^*\**^(*T* − 1). With each subsequent restart of the optimization scheme at *T* as described in Appendix C, we sample ξ_*i*_, *i* ∈ {1,…, 30}, from a uniform distribution 𝒰 [0, 0.01]. We then clamp to 0 any fibrillar activation 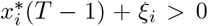, *i* ∈ {1,…, 28}, to ensure a contractile initial guess.

## Appendix C. Choice of optimization schemes

We found different optimization schemes to exhibit varying degrees of performance across the evaluated motions and their constituent phases. Here, we provide a list of the optimization schemes that we utilize throughout the study. In the list below, we employ the algorithm naming convention from the Julia Optimization.jl package [66].

- Main optimization problems:
  – Global optimization:
    ∗ `NOMADOpt:` Mesh Adaptive Direct Search algorithm (MADS) for blackbox optimization [67] from the NOMAD.jl package [68].
  – Local optimization:
    ∗ `LN_SBPLX: Sblx` algorithm from the NLopt library [70] based on the Subplex algorithm [69].
- Auxiliary optimization problems:
  – Global optimization:
    ∗ `G_MLSL_LDS:` Multi-Level Single-Linkage (MLSL) multistart algorithm [71, 72] using Low-Discrepancy Sequences (LDS) [73] from the NLopt library [70] together with with the Nelder-Mead method [74] as the local optimization scheme LN NELDERMEAD from NLopt [70].
    ∗ `BBO_adaptive_de_rand_1_bin_radiuslimited`: multi-threaded adaptive differential evolution optimizer with radius-limited sampling from the Julia blackbox optimization package BlackBoxOptim.jl [75].

We apply these optimization schemes to compute the different motion phases as follows:

- Picking a fruit:
  – First and third phases:
    ∗ Main optimization problems: local minimization using `LN_SBPLX.`
    ∗ Auxiliary optimization problems: global minimization using `G_MLSL_LDS` with the LN_NELDERMEAD local optimization scheme.
  – Second phase:
    ∗ Main optimization problems: local minimization using `LN_SBPLX.`
    ∗ Auxiliary optimization problems: not applicable.
- Lifting a log:
  – All three phases:
    ∗ Main optimization problems: local minimization using `LN_SBPLX.`
    ∗ Auxiliary optimization problems: global minimization using `BBO_adaptive_de_rand_1_ bin_radiuslimited.`
- Asymmetric lifting of a log:
  – First phase: solution **x**^*\**^(1) obtained from the symmetric lifting scenario.
  – Second and third phases:
    ∗ Main optimization problems: global minimization using `NOMADOpt`. We replaced `LN_SBPLX.` with the `NOMADOpt` global optimization scheme since minimizing locally around auxiliary optima was no longer a viable approach for the asymmetric lifting case.
    ∗ Auxiliary optimization problems: not applicable.

## References

[1] W. M. Kier, K. K. Smith, Tongues, tentacles and trunks: The biomechanics of movement in muscular-hydrostats, Zoological Journal of the Linnean Society 83 (4) (1985) 307–324. doi:10.1111/j.1096-3642.1985.tb01178.x.

[2] W. M. Kier, M. P. Stella, The arrangement and function of octopus arm musculature and connective tissue, Journal of Morphology 268 (10) (2007) 831–843. doi:10.1002/jmor.10548.

[3] W. M. Kier, The diversity of hydrostatic skeletons, Journal of Experimental Biology 215 (8) (2012) 1247–1257. doi:10.1242/jeb.056549.

[4] L. L. Longren, L. Eigen, A. Shubitidze, O. Lieschnegg, D. Baum, J. A. Nyakatura, T. Hildebrandt, M. Brecht, Dense reconstruction of elephant trunk musculature, Current Biology 33 (21) (2023) 4713–4720.e3. doi:10.1016/j.cub.2023.09.007.

[5] J. Wu, Y. Zhao, Y. Zhang, D. Shumate, S. Braccini Slade, S. V. Franklin, D. L. Hu, Elephant trunks form joints to squeeze together small objects, Journal of The Royal Society Interface 15 (147) (2018) 20180377. doi:10.1098/rsif.2018.0377.

[6] R. Cornette, A. Delapré, C. Houssin, B. Mulot, E. Pouydebat, Measuring the force of the tip of the elephants trunk, MethodsX 9 (2022) 101896. doi:10.1016/j.mex.2022.101896.

[7] J. Shoshani, Understanding proboscidean evolution: A formidable task, Trends in Ecology & Evolution 13 (12) (1998) 480–487. doi:10.1016/S0169-5347(98)01491-8.

[8] G. Kirchhoff, Ueber das Gleichgewicht und die Bewegung eines unendlich dünnen elastischen Stabes., Journal für die reine und angewandte Mathematik 1859 (56) (1859) 285–313. doi: 10.1515/crll.1859.56.285.

[9] A. E. H. Love, A Treatise on the Mathematical Theory of Elasticity, 2nd Edition, at the University Press, Cambridge, 1906.

[10] S. S. Antman, The Theory of Rods, in: C. Truesdell (Ed.), Linear Theories of Elasticity and Thermoelasticity: Linear and Nonlinear Theories of Rods, Plates, and Shells, Springer, Berlin, Heidelberg, 1973, pp. 641–703. doi:10.1007/978-3-662-39776-3_6.

[11] J. C. Simo, A finite strain beam formulation. The three-dimensional dynamic problem. Part I, Computer Methods in Applied Mechanics and Engineering 49 (1) (1985) 55–70. doi:10.1016/0045-7825(85)90050-7.

[12] E. H. Dill, Kirchhoff’s Theory of Rods, Archive for History of Exact Sciences 44 (1) (1992) 1–23. 41133926.

[13] M. Bergou, M. Wardetzky, S. Robinson, B. Audoly, E. Grinspun, Discrete elastic rods, in: ACM SIGGRAPH 2008 Papers, SIGGRAPH ‘08, Association for Computing Machinery, New York, NY, USA, 2008, pp. 1–12. doi:10.1145/1399504.1360662.

[14] H. Altenbach, V. A. Eremeyev, F. Pfeiffer, F. G. Rammerstorfer, J. Salençon, B. Schrefler, P. Serafini (Eds.), Generalized Continua from the Theory to Engineering Applications, Vol. 541 of CISM International Centre for Mechanical Sciences, Springer, Vienna, 2013. doi: 10.1007/978-3-7091-1371-4.

[15] A. Lazarus, J. T. Miller, P. M. Reis, Continuation of equilibria and stability of slender elastic rods using an asymptotic numerical method, Journal of the Mechanics and Physics of Solids 61 (8) (2013) 1712–1736. doi:10.1016/j.jmps.2013.04.002.

[16] M. Gazzola, L. H. Dudte, A. G. McCormick, L. Mahadevan, Forward and inverse problems in the mechanics of soft filaments, Royal Society Open Science 5 (6) (2018) 171628. doi: 10.1098/rsos.171628.

[17] M. Hubbard, Dynamics of the pole vault, Journal of Biomechanics 13 (11) (1980) 965–976. doi:10.1016/0021-9290(80)90168-2.

[18] P. Gudmundson, The dynamic behaviour of slender structures with cross-sectional cracks, Journal of the Mechanics and Physics of Solids 31 (4) (1983) 329–345. doi:10.1016/0022-5096(83)90003-0.

[19] G. H. M. van der Heijden, S. Neukirch, V. G. A. Goss, J. M. T. Thompson, Instability and self-contact phenomena in the writhing of clamped rods, International Journal of Mechanical Sciences 45 (1) (2003) 161–196. doi:10.1016/S0020-7403(02)00183-2.

[20] S. Neukirch, B. Roman, B. de Gaudemaris, J. Bico, Piercing a liquid surface with an elastic rod: Buckling under capillary forces, Journal of the Mechanics and Physics of Solids 55 (6) (2007) 1212–1235. doi:10.1016/j.jmps.2006.11.009.

[21] G.-F. Wang, X.-Q. Feng, Timoshenko beam model for buckling and vibration of nanowires with surface effects, Journal of Physics D: Applied Physics 42 (15) (2009) 155411. doi: 10.1088/0022-3727/42/15/155411.

[22] K. Chandraseker, S. Mukherjee, J. T. Paci, G. C. Schatz, An atomistic-continuum Cosserat rod model of carbon nanotubes, Journal of the Mechanics and Physics of Solids 57 (6) (2009) 932–958. doi:10.1016/j.jmps.2009.02.005.

[23] T. Bretl, Z. McCarthy, Quasi-static manipulation of a Kirchhoff elastic rod based on a geometric analysis of equilibrium configurations, The International Journal of Robotics Research 33 (1) (2014) 48–68. doi:10.1177/0278364912473169.

[24] W. Wu, X. Cao, Mechanics model and its equation of wire rope based on elastic thin rod theory, International Journal of Solids and Structures 102–103 (2016) 21–29. doi:10.1016/j.ijsolstr.2016.10.021.

[25] N. Lv, J. Liu, H. Xia, J. Ma, X. Yang, A review of techniques for modeling flexible cables, Computer-Aided Design 122 (2020) 102826. doi:10.1016/j.cad.2020.102826.

[26] W. Lacarbonara, H. Yabuno, Refined models of elastic beams undergoing large in-plane motions: Theory and experiment, International Journal of Solids and Structures 43 (17) (2006) 5066–5084. doi:10.1016/j.ijsolstr.2005.07.018.

[27] M. K. Jawed, F. Da, J. Joo, E. Grinspun, P. M. Reis, Coiling of elastic rods on rigid substrates, Proceedings of the National Academy of Sciences 111 (41) (2014) 14663–14668. doi:10.1073/pnas.1409118111.

[28] T. Yu, J. A. Hanna, Bifurcations of buckled, clamped anisotropic rods and thin bands under lateral end translations, Journal of the Mechanics and Physics of Solids 122 (2019) 657–685. doi:10.1016/j.jmps.2018.01.015.

[29] V. A. Aloi, D. C. Rucker, Estimating Loads Along Elastic Rods, in: 2019 International Conference on Robotics and Automation (ICRA), 2019-05-20/2019-05-24, pp. 2867–2873. doi:10.1109/ICRA.2019.8794301.

[30] B. Chakrabarti, Y. Liu, J. LaGrone, R. Cortez, L. Fauci, O. du Roure, D. Saintillan, A. Lindner, Flexible filaments buckle into helicoidal shapes in strong compressional flows, Nature Physics 16 (6) (2020) 689–694. doi:10.1038/s41567-020-0843-7.

[31] A. Sintov, S. Macenski, A. Borum, T. Bretl, Motion Planning for Dual-Arm Manipulation of Elastic Rods, IEEE Robotics and Automation Letters 5 (4) (2020) 6065–6072. doi:10.1109/LRA.2020.3011352.

[32] P. Johanns, P. Grandgeorge, C. Baek, T. G. Sano, J. H. Maddocks, P. M. Reis, The shapes of physical trefoil knots, Extreme Mechanics Letters 43 (2021) 101172. doi:10.1016/j.eml.2021.101172.

[33] T. G. Sano, M. Pezzulla, P. M. Reis, A Kirchhoff-like theory for hard magnetic rods under geometrically nonlinear deformation in three dimensions, Journal of the Mechanics and Physics of Solids 160 (2022) 104739. doi:10.1016/j.jmps.2021.104739.

[34] W. Huang, X. Huang, C. Majidi, M. K. Jawed, Dynamic simulation of articulated soft robots, Nature Communications 11 (1) (2020) 2233. doi:10.1038/s41467-020-15651-9.

[35] D. Trivedi, A. Lotfi, C. D. Rahn, Geometrically Exact Models for Soft Robotic Manipulators, IEEE Transactions on Robotics 24 (4) (2008) 773–780. doi:10.1109/TRO.2008.924923.

[36] L. Wang, Y. Kim, C. F. Guo, X. Zhao, Hard-magnetic elastica, Journal of the Mechanics and Physics of Solids 142 (2020) 104045. doi:10.1016/j.jmps.2020.104045.

[37] N. Naughton, J. Sun, A. Tekinalp, T. Parthasarathy, G. Chowdhary, M. Gazzola, Elastica: A Compliant Mechanics Environment for Soft Robotic Control, IEEE Robotics and Automation Letters 6 (2) (2021) 3389–3396. doi:10.1109/LRA.2021.3063698.

[38] P. Dagenais, S. Hensman, V. Haechler, M. C. Milinkovitch, Elephants evolved strategies reducing the biomechanical complexity of their trunk, Current Biology 31 (21) (2021) 4727–4737.e4. doi:10.1016/j.cub.2021.08.029.

[39] A. K. Schulz, M. Boyle, C. Boyle, S. Sordilla, C. Rincon, S. Hooper, C. Aubuchon, J. S. Reidenberg, C. Higgins, D. L. Hu, Skin wrinkles and folds enable asymmetric stretch in the elephant trunk, Proceedings of the National Academy of Sciences 119 (31) (2022) e2122563119. doi:10.1073/pnas.2122563119.

[40] L. Purkart, J. M. Tuff, M. Shah, L. V. Kaufmann, C. Altringer, E. Maier, U. Schneeweiß, E. Tunckol, L. Eigen, S. Holtze, G. Fritsch, T. Hildebrandt, M. Brecht, Trigeminal ganglion and sensory nerves suggest tactile specialization of elephants, Current Biology 32 (4) (2022) 904–910.e3. doi:10.1016/j.cub.2021.12.051.

[41] L. V. Kaufmann, U. Schneeweiß, E. Maier, T. Hildebrandt, M. Brecht, Elephant facial motor control, Science Advances 8 (43) (2022) eabq2789. doi:10.1126/sciadv.abq2789.

[42] N. Deiringer, U. Schneeweiß, L. V. Kaufmann, L. Eigen, C. Speissegger, B. Gerhardt, S. Holtze, G. Fritsch, F. Göritz, R. Becker, A. Ochs, T. Hildebrandt, M. Brecht, The functional anatomy of elephant trunk whiskers, Communications Biology 6 (1) (2023) 1–12. doi:10.1038/s42003-023-04945-5.

[43] J. F. Wilson, U. Mahajan, S. A. Wainwright, L. J. Croner, A continuum model of elephant trunks, Journal of Biomechanical Engineering 113 (1) (1991) 79–84. doi:10.1115/1.2894088.

[44] Y. Liang, R. M. McMeeking, A. G. Evans, A finite element simulation scheme for biological muscular hydrostats, Journal of Theoretical Biology 242 (1) (2006) 142–150. doi:10.1016/j.jtbi.2006.02.008.

[45] A. K. Schulz, J. S. Reidenberg, J. N. Wu, C. Y. Tang, B. Seleb, J. Mancebo, N. Elgart, D. L. Hu, Elephant trunks use an adaptable prehensile grip, Bioinspiration & Biomimetics 18 (2) (2023) 026008. doi:10.1088/1748-3190/acb477.

[46] B. Kaczmarski, S. Leanza, R. Zhao, E. Kuhl, D. E. Moulton, A. Goriely, Minimal Design of the Elephant Trunk as an Active Filament, Physical Review Letters 132 (24) (2024) 248402. doi:10.1103/PhysRevLett.132.248402.

[47] B. Kamare, M. L. Preti, I. Bernardeschi, S. Lantean, P. Dagenais, M. Milinkovitch, L. Beccai, Study and Preliminary Modeling of Microstructure and Morphology of the Elephant Trunk Skin, in: F. Meder, A. Hunt, L. Margheri, A. Mura, B. Mazzolai (Eds.), Biomimetic and Biohybrid Systems, Springer Nature Switzerland, Cham, 2023, pp. 101–114. doi:10.1007/978-3-031-39504-8_7.

[48] D. Trivedi, C. D. Rahn, W. M. Kier, I. D. Walker, Soft Robotics: Biological Inspiration, State of the Art, and Future Research, Applied Bionics and Biomechanics 5 (3) (2008) 99–117. doi:10.1080/11762320802557865.

[49] R. Ciéslak, A. Morecki, Elephant trunk type elastic manipulator - a tool for bulk and liquid materials transportation, Robotica 17 (1) (1999) 11–16. doi:10.1017/S0263574799001009.

[50] M. W. Hannan, I. D. Walker, Kinematics and the Implementation of an Elephant’s Trunk Manipulator and Other Continuum Style Robots, Journal of Robotic Systems 20 (2) (2003) 45–63. doi:10.1002/rob.10070.

[51] M. Rolf, J. J. Steil, Efficient Exploratory Learning of Inverse Kinematics on a Bionic Elephant Trunk, IEEE Transactions on Neural Networks and Learning Systems 25 (6) (2014) 1147–1160. doi:10.1109/TNNLS.2013.2287890.

[52] C. Laschi, B. Mazzolai, M. Cianchetti, Soft robotics: Technologies and systems pushing the boundaries of robot abilities, Science Robotics 1 (1) (2016) eaah3690. doi:10.1126/scirobotics.aah3690.

[53] A. Tekinalp, N. Naughton, S. H. Kim, U. Halder, R. Gillette, P. G. Mehta, W. Kier, M. Gazzola, Topology, dynamics, and control of a muscle-architected soft arm, Proceedings of the National Academy of Sciences 121 (41) (2024) e2318769121. doi:10.1073/pnas.2318769121.

[54] B. Kaczmarski, D. E. Moulton, E. Kuhl, A. Goriely, Active filaments I: Curvature and torsion generation, Journal of the Mechanics and Physics of Solids 164 (2022) 104918. doi:10.1016/j.jmps.2022.104918.

[55] A. Goriely, The Mathematics and Mechanics of Biological Growth, Springer Verlag, New York, 2017.

[56] D. E. Moulton, T. Lessinnes, A. Goriely, Morphoelastic rods. Part I: A single growing elastic rod, Journal of the Mechanics and Physics of Solids 61 (2) (2013) 398–427. doi:10.1016/j.jmps.2012.09.017.

[57] S. Xu, X. Yue, C. Zhang, Homogenization: In mathematics or physics?, Discrete and Continuous Dynamical Systems - S 9 (5) (2016) 1575–1590. doi:10.3934/dcdss.2016064.

[58] D. E. Moulton, T. Lessinnes, A. Goriely, Morphoelastic rods III: Differential growth and curvature generation in elastic filaments, Journal of the Mechanics and Physics of Solids 142 (2020) 104022. doi:10.1016/j.jmps.2020.104022.

[59] S. Leanza, J. Lu-Yang, B. Kaczmarski, S. Wu, E. Kuhl, R. R. Zhao, Elephant Trunk Inspired Multimodal Deformations and Movements of Soft Robotic Arms, Advanced Functional Materials 34 (29) (2024) 2400396. doi:10.1002/adfm.202400396.

[60] U. M. Ascher, R. M. M. Mattheij, R. D. Russell, Appendix B: A Collocation Code, in: Numerical Solution of Boundary Value Problems for Ordinary Differential Equations, Classics in Applied Mathematics, Society for Industrial and Applied Mathematics, 1995, pp. 526–533. doi:10.1137/1.9781611971231.appb.

[61] J. Méndez, Density and composition of mammalian muscle., Metabolism 9 (1960) 184–188.

[62] G. K. Klute, J. M. Czerniecki, B. Hannaford, McKibben artificial muscles: Pneumatic actuators with biomechanical intelligence, in: 1999 IEEE/ASME International Conference on Advanced Intelligent Mechatronics (Cat. No.99TH8399), 0019/1999-09-23, pp. 221–226. doi:10.1109/AIM.1999.803170.

[63] B. Kaczmarski, A. Goriely, E. Kuhl, D. E. Moulton, A Simulation Tool for Physics-Informed Control of Biomimetic Soft Robotic Arms, IEEE Robotics and Automation Letters 8 (2) (2023) 936–943. doi:10.1109/LRA.2023.3234819.

[64] B. Kaczmarski, D. E. Moulton, A. Goriely, E. Kuhl, Bayesian design optimization of biomimetic soft actuators, Computer Methods in Applied Mechanics and Engineering 408 (2023) 115939. doi:10.1016/j.cma.2023.115939.

[65] B. Kaczmarski, D. E. Moulton, A. Goriely, E. Kuhl, Minimal activation with maximal reach: Reachability clouds of bio-inspired slender manipulators, Extreme Mechanics Letters 71 (2024) 102207. doi:10.1016/j.eml.2024.102207.

[66] V. K. Dixit, C. Rackauckas, Optimization.jl: A unified optimization package, Zenodo (2023). doi:10.5281/zenodo.7738525.

[67] C. Audet, S. Le Digabel, V. Rochon Montplaisir, C. Tribes, Algorithm 1027: NOMAD version 4: Nonlinear optimization with the MADS algorithm, ACM Transactions on Mathematical Software 48 (3) (2022) 35:1–35:22. doi:10.1145/3544489.

[68] A. Montoison, P. Pascal, L. Salomon, NOMAD.jl: A Julia interface for the constrained black-box solver NOMAD (Jul. 2020). doi:10.5281/zenodo.3700167.

[69] T. H. Rowan, Functional stability analysis of numerical algorithms, Ph.D. thesis, University of Texas at Austin, USA (1990).

[70] S. G. Johnson, The NLopt nonlinear-optimization package, https://github.com/stevengj/nlopt (2007).

[71] A. H. G. Rinnooy Kan, G. T. Timmer, Stochastic global optimization methods part I: Clustering methods, Mathematical Programming 39 (1) (1987) 27–56. doi:10.1007/BF02592070.

[72] A. H. G. Rinnooy Kan, G. T. Timmer, Stochastic global optimization methods part II: Multi level methods, Mathematical Programming 39 (1) (1987) 57–78. doi:10.1007/BF02592071.

[73] S. Kucherenko, Y. Sytsko, Application of Deterministic Low-Discrepancy Sequences in Global Optimization, Computational Optimization and Applications 30 (3) (2005) 297–318. doi: 10.1007/s10589-005-4615-1.

[74] J. A. Nelder, R. Mead, A Simplex Method for Function Minimization, The Computer Journal 7 (4) (1965) 308–313. doi:10.1093/comjnl/7.4.308.

[75] R. Feldt, BlackBoxOptim.jl, https://github.com/robertfeldt/BlackBoxOptim.jl (2018).

